# Spatial and Multiomic profiling of muscle regeneration dynamics in Duchenne Muscular Dystrophy

**DOI:** 10.64898/2025.12.02.691824

**Authors:** L. Virtanen, C. D’Ercole, L. Saillard, F. Grandi, C. Peccate, B. Hoareau, V Karunanuthy, C Bertholle, B. Izac, B Saintpierre, M Andrieu, F Letourneur, K Mamchaoui, M Chapart, S. Vasseur, P. Smeriglio, T Evangelista, L Madaro, L. Giordani

## Abstract

Duchenne muscular dystrophy (DMD) is a pediatric degenerative myopathy caused by the absence of functional dystrophin. As a result, DMD muscles exhibit compromised myofiber integrity and increased susceptibility to mechanical damage. In early disease stages, muscles undergo repeated cycles of degeneration and regeneration; over time, however, this regenerative capacity declines, leading to the gradual replacement of muscle tissue with fat and fibrosis. While several signaling pathways have been identified as deregulated in dystrophic muscle, the cellular and molecular mechanisms underlying this regenerative exhaustion remain to be fully elucidated. To address this, we constructed a comprehensive cellular atlas of human dystrophic muscle using high-resolution spatial transcriptomics (Visium HD), capturing the cellular crosstalk within regenerative regions.

Cell–to-cell communication analysis revealed activation of Notch signaling mediated by NOTCH3 in activated satellite cells. Immunostaining confirmed elevated NOTCH3 expression in both DMD patient samples and in the mdx mouse model at late disease stages. Silencing of NOTCH3 in primary myoblasts improved myogenic differentiation, pinpointing NOTCH3-mediated signaling as a contributor to regeneration impairment. To further dissect the dynamics of regeneration and infer the gene regulatory networks governing myogenic differentiation, we integrated paired snRNA-seq and snATAC-seq data from young, adult, and aged mdx mice. This analysis identified GLIS3 upregulation as an additional barrier to effective myogenesis. GLIS3 displayed an overall increase in dystrophic muscles, while silencing experiments enhanced differentiation in myoblasts.

Together, our work reveals intrinsic defects in the dystrophic stem cell compartment that emerge during disease progression and hinder the execution of the myogenic program. These findings suggest NOTCH3 and GLIS3 as potential therapeutic targets to enhance regeneration and maintain muscle integrity in DMD. This study provides a high-resolution map of the dystrophic regenerative landscape and offers a valuable resource for future translational research.

## Introduction

Duchenne muscular dystrophy (DMD) is an X-linked recessive disorder that primarily affects male children. Symptoms typically emerge between 3 and 6 years of age, with patients showing delayed mobility and calf pseudohypertrophy. Muscle weakness progresses from the legs and pelvis to the trunk, leading to loss of ambulation, respiratory impairment, cardiac complications, and reduced lifespan^1,2^.

Skeletal muscles in DMD patients exhibit two major differences from healthy muscle. The first is muscle fiber fragility, such that mild mechanical stress leads to muscle damage, which remains constant throughout disease progression; the second is impaired repair capacity, which develops over time. In DMD, muscle regeneration differs significantly from that in healthy muscle due to the chronic nature of the disease. Unlike healthy muscle, where regeneration is acute and efficient, DMD muscle is in a constant state of repair in a pathological environment characterized by chronic inflammation and fibrosis^3–5^, which impair regeneration and promote tissue scarring instead of functional muscle repair. Multiple lines of evidence, primarily from mouse models, suggest that disrupted cellular communication^6^ and intrinsic defects in the stem cell compartment could contribute to the pathogenesis of the disease once the initial fiber damage occurs^7^. Specifically, the asynchronous and focal nature of regeneration hinders repair by providing conflicting signaling to the regenerating areas^3^. The chronically inflamed environment perturbs the crosstalk between fibro-adipogenic progenitors (FAPs) and macrophages, driving FAPs toward a matrix-producing, profibrotic phenotype^6^. Consistently, several extracellular matrix components have been reported to be differentially expressed between healthy and dystrophic FAPs^8^. In parallel, the absence of dystrophin disrupts stem cell polarity from the earliest stages of disease, contributing to reduced regenerative efficiency^7^. In addition, multiple signaling axis—including STAT3^9^, p38^10^, and NOTCH^11,12^—have been shown to be deregulated in dystrophic MuSCs. For these reasons cellular heterogeneity in DMD has been widely investigated using single-cell RNA sequencing (scRNA-seq) and, more recently, spatial transcriptomics in mouse models, higher primates and human patients^8,13–16^. These studies have been focused mainly on differences between healthy and diseased muscle characterizing the main features of the disease; however, the molecular drivers of the loss of regenerative potential remain to be fully elucidated. This is partially due to the intrinsic challenge of comparing inherently different tissues: a healthy, non-regenerating muscle versus dystrophic muscle in which both degeneration and regeneration are actively ongoing. Moreover, given the presence of multiple asynchronous regenerating foci within the dystrophic muscle^3^, even classic regenerative models such as notexin injection, in which the whole muscle is subjected to injury, present several limitations as a direct reference for regeneration in DMD^17^. Indeed, without the ability to analyze individual regenerative foci within dystrophic muscle, identifying specific defects remains a significant challenge.

To bridge this gap, we chose to directly investigate regeneration within dystrophic muscle using two complementary strategies (Fig. 1 A). First, we applied high-resolution spatial transcriptomics to human muscle biopsies from DMD patients undergoing surgical treatment. This allowed us to identify regenerative foci and characterize local cellular crosstalk and signaling activity across different stages, from activated satellite cells to mature fibers. Notably, we identified an activation of Notch signaling mediated via NOTCH3 within the satellite cell compartment, which resulted in less efficient myogenic differentiation. In parallel, to characterize the different regulatory networks controlling the myogenic program, we extended our study to the classical mdx dystrophic mouse model. We performed paired snRNA-seq and snATAC-seq (single-nucleus Multiome) at multiple time points (2 weeks, 5 weeks, 6 months, and 14 months). This enabled us to follow disease progression and identify how changes in transcription factors alter the normal myogenic trajectory. This analysis highlighted the progressive upregulation of *Glis3* (GLIS Family Zinc Finger 3) as a potential barrier to efficient MuSCs (Muscle Stem Cells) activation, contributing to a reduced pool of committed progenitors. Collectively, our findings identified the progressive deregulation of Notch signaling and the upregulation of Glis3 in muscle stem cells as potential contributors to impaired regeneration in dystrophic muscle. These insights lay the groundwork for future therapeutic strategies aimed at restoring and supporting effective regeneration in dystrophic muscle.

**Figure 1.**
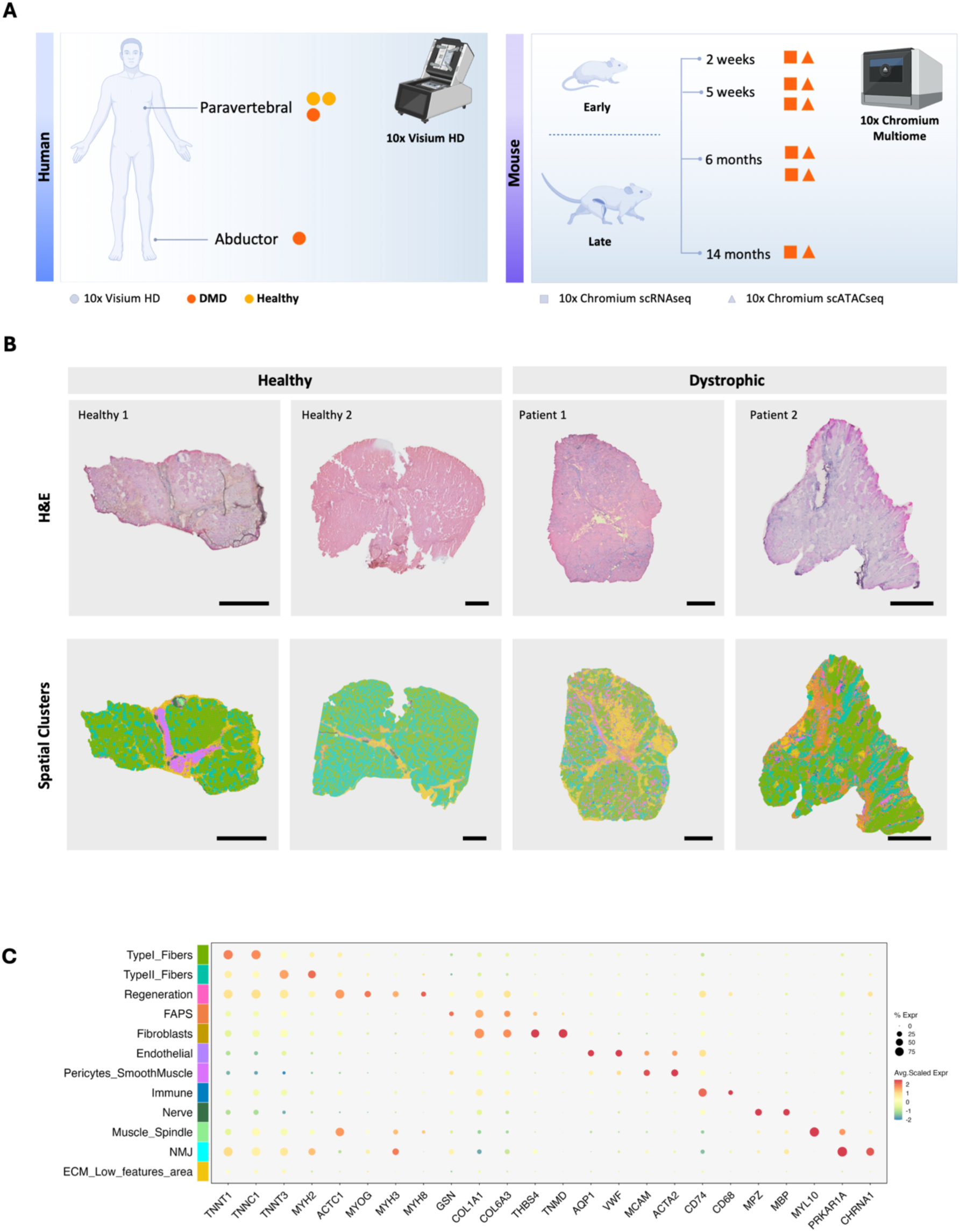
Identification of different cell populations in human skeletal muscle using high-resolution spatial transcriptomics. A) Diagram illustrating the workflow and experimental strategy. Healthy human (n=2) and DMD patient (n=2) muscle biopsies were used for the HD spatial transcriptomics. Hindlimb muscles from two C57BL/10-*mdx* mice per age group (2 weeks, 5weeks, 6months and 14 months) were pooled for combined snRNA and snATAC-seq. B) H&E image and spatial mapping of four Visium samples (CTR1 Healthy, CTR2 Healthy, DMD1 patient, DMD2 patient) with 8 μm bins colored on the basis of unsupervised clustering. Scale bar, 1 mm. C) Dot plot showing the expression values of canonical gene markers for each cluster identified in the spatial human dataset.

## Results

### High-resolution spatial transcriptomics accurately captures regeneration process within DMD muscle

To study spatial gene expression patterns within the regenerative environment in the dystrophic muscle, we performed high-resolution spatial transcriptomics (Visium HD – 10x Genomics) on a set of dystrophic muscles and healthy controls obtained from individuals undergoing surgical interventions displaying different degrees of regeneration and fibrosis (2 dystrophic and 2 healthy controls, aged between 14 and 17 years – Extended Data Fig. 1A and Supplementary Table 1). In brief, frozen muscle tissues were sectioned onto Visium HD slides, stained with hematoxylin and eosin (H&E), imaged, and processed to capture barcoded mRNA, which was then mapped back to the tissue section^18^. Data was processed with Space Ranger and analyzed in Seurat 5^19^ using aggregated 8 µm bins (∼1,000,000 bins, averaging 111 features each). In our initial analysis we observed marked regional variability in transcript density, which correlated with different tissue regions and the preservation status of the analyzed muscle samples (Extended Data Fig. 1B-C). Notably, collagen-enriched areas presented a lower transcript density than regions containing muscle fibers or cellular infiltrates. Furthermore, specific areas presented limited transcript detection, likely due to local mRNA degradation; these two regions, clustered together, were annotated as ECM/low-feature areas (Fig. 1B-C and Extended Data Fig. 1C).

To cluster and annotate the different cell populations and functional domains in our dataset, we first applied a low-resolution clustering approach identifying five major groups – roughly corresponding to fibers, mesenchymal cells, endothelial and other vessel-associated cells, immune cells, and nerves. Subsequently we analyzed each group separately to resolve specific populations. Consistent with previous studies utilizing Visium technology in murine muscle^20–24^, we classified the bins into ten major categories: Type I and Type II muscle fibers, fibro-adipogenic progenitors (FAPs), fibroblasts, endothelial cells, pericytes and smooth muscle cells, immune cells, nerves and regeneration-associated bins (Fig. 1B and C). Additionally, we were able to identify muscle spindle regions on the basis of their morphological location and expression of *MYL10*^25^ (Fig. 1C) for a total of eleven clusters. To correctly label the regenerating regions, which typically consist of multiple distinct cell types, we used the expression of canonical markers associated with myogenic progression such as *MYOG*, *MYH3*, *MYH8*, and *ACTC1* which was recently identified as a marker of a fiber subpopulation supporting MuSCs during regeneration^26^ (Fig. 1C).

One of the key advantages of spatial transcriptomics is the ability to cross-validate the quality of annotations with morphological features. By directly inspecting the samples, we confirmed that among our annotated populations, those recognizable via H&E staining aligned accurately with their expected anatomical compartments (Extended Data Fig. 2A). For clusters such as Type I and Type II muscle fibers, which cannot be distinguished solely on the basis of H&E staining, we utilized a module score derived from a set of fiber type-specific genes^27,28^ and validated these results through immunofluorescence staining for specific fiber type markers in serial sections (Fig. 2A and Extended Data Fig. 2B). For NMJ regions, we performed Bungarotoxin staining on consecutive sections confirming enrichment in the predicted areas (Extended Data Fig. 2C). As expected, areas associated with mesenchymal cells (FAPs and fibroblasts), immune infiltration, and regeneration were predominantly present in the DMD samples (Fig. 1B and Supplementary Table 2). Notably, the regenerative area accounted for up to 9% of the muscle area in DMD samples but only ∼0.05% in healthy controls (Fig. 2B and Supplementary Table 2). ECM and low-feature areas were also more prevalent in DMD samples, which is consistent with the progression of fibrosis but may also reflect variations in sample quality.

**Figure 2.**
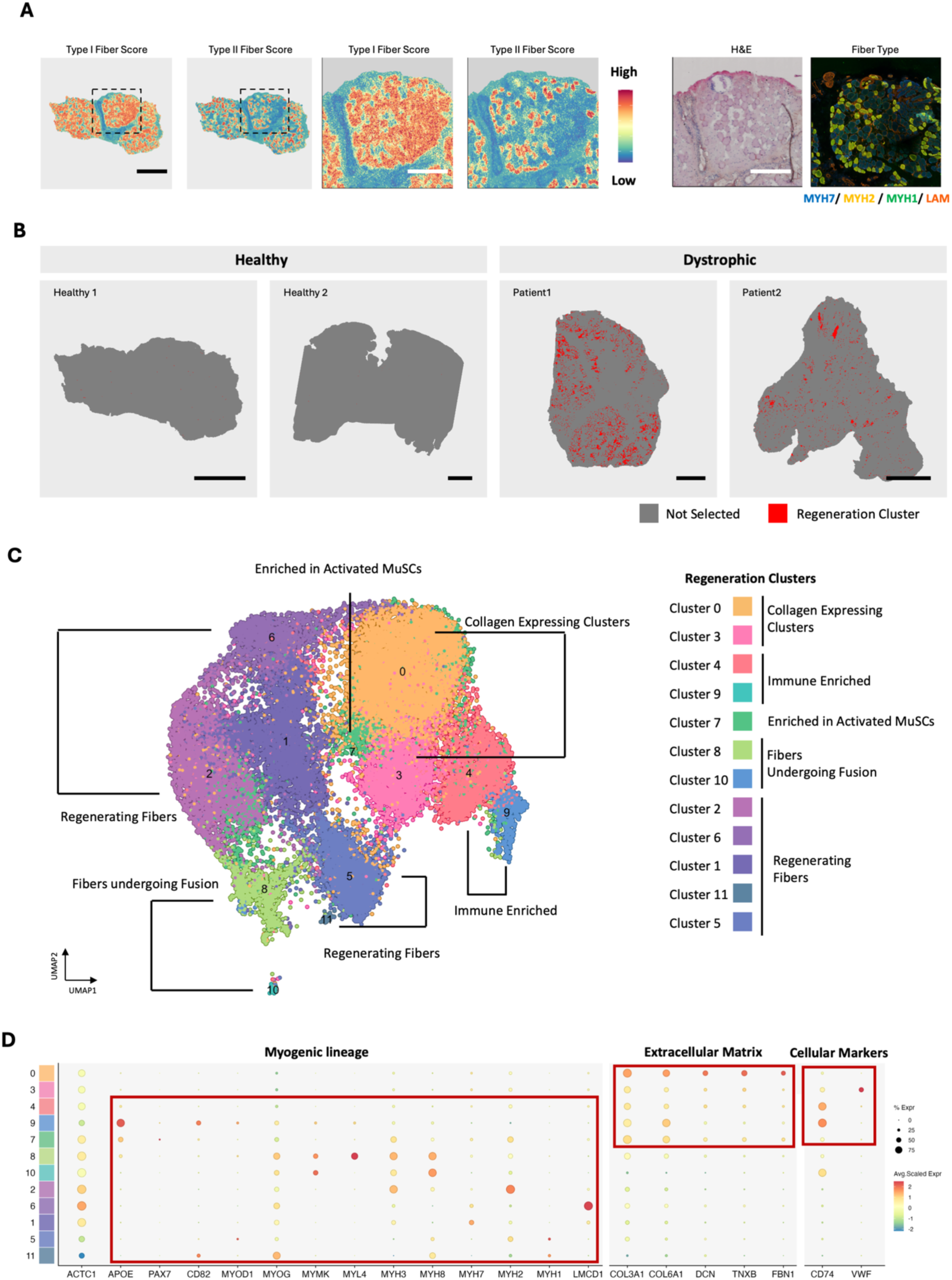
– Subclustering of regeneration cluster identifies myogenic differentiation continuum from patient muscle biopsies. A) Spatial distribution and relative expression levels of the fiber type score in CTR1. Scale bar: 1 mm. Score for type I and type II fibers in the region highlighted by the dashed box (left). Scale bar: 200 µm. H&E staining and validation of the same region (right). To validate the fiber score, serial sections were immunostained for Myh7 (blue), Myh2 (yellow), Myh1 (green), and Laminin (red). Scale bar: 200 µm. B) Visualization of regenerative cluster spatial distribution in healthy controls and dystrophic patients. Bins assigned to regenerative areas are highlighted in red. Scale bar, 1 mm. C) Uniform manifold approximation and projection (UMAP) visualization and subclustering of the regeneration clusters; spots are color-coded and labeled on the basis of subcluster assignment. D) Dot plot showing the expression values of selected genes across different clusters within the regenerating continuum. Red boxes highlight distinct gene sets within specific clusters.

After annotating the different tissue regions, we aimed to characterize the different stages along the regeneration continuum by subclustering the regeneration areas. We obtained twelve different clusters (Fig. 2C). Differences in the expression of PAX7, *MYOG, MYH3, MYH8*, and mature fiber markers such as *MYH2*, MYH1 and MYH7 clearly indicate differentiation progression across the clusters (Fig. 2D). Among the identified clusters, Cluster 7 was enriched with activated MuSC and characterized by the expression of *PAX7* (∼10% of the bins), *APOE* and *MYOG*. Early myogenic stages were captured by clusters 8 and 10, representing fibers undergoing active myonuclear fusion, as indicated by the expression of *MYMK.* Cluster 2 displayed expression of *MYOG, MYH3,* and low levels of *MYH8* but showed almost no *MYMK* expression, suggesting a more advanced stage. Clusters 6 and 1 expressed *MYOG* together with *LMCD1*, a gene linked to skeletal muscle hypertrophy and regrowth^29^, while showing low levels of MYH3 and the type I fiber marker MYH7, consistent with an intermediate differentiation state. In contrast, clusters 5 and 11 displayed minimal *LMCD1* expression but expressed *MYH2*, *MYH1*, and the regenerative fiber marker *MYH8*, characteristic of later stages of myofiber maturation. Beyond the myogenic core, we identified two interstitial clusters (0 and 3) strongly expressing *COL3A1* and *COL6A1*, which are collagens that are rapidly upregulated during regeneration^30,31^. These clusters also expressed *TNXB* and *FBN1*, similarly to CD55+ *FAP*s population previously identified by Fitzgerald^32^. These clusters also exhibited low levels of *MYOG* and *MYH3* expression, suggesting that they may represent a mix of mesenchymal and myogenic cells surrounding regenerating fibers in early stages. Additionally, the expression of *VWF* in cluster 3, was consistent with its interstitial localization and suggests the presence of endothelial cells within or adjacent to this cluster (Fig. 2D). Intriguingly, clusters 4 and 9 showed strong expression of *CD74*, indicating an enrichment of immune cells within these populations. Notably, cluster 9 also exhibited moderate expression of myogenic markers, including *MYOG*, *MYOD*, and *MYMK*, suggesting that it may represent an interface region where immune and myogenic compartments interact.

In summary, these results show that high-resolution spatial transcriptomics can accurately identify specific cell populations in human muscle, as confirmed by H&E and immunofluorescence staining. Moreover, through unbiased iterative subclustering of regenerative regions, we were able to resolve and characterize distinct stages along the myogenic differentiation continuum.

### Cell–cell communication mapping identifies local signals driving regeneration

As our results above establish that ST accurately capture the regeneration process in human DMD muscle, we next investigated the cellular communication network within the regenerating areas to better understand how the dystrophic microenvironment influences the myogenic program. Specifically, we focused our analysis on the set of clusters expressing myogenic lineage genes, aiming to capture interactions across different stages of regeneration—including early-stage fibers undergoing myoblast fusion (Cluster 8), more developed fibers (Clusters 2), immune enriched (Cluster 9), and activated MuSCs (Cluster 7). To gain insight into the cellular crosstalk mediated by these clusters in the regenerating areas, we selected three representative regions enriched in fusing myoblasts, immune infiltration, or activated satellite cells and performed ligand‒receptor analysis using the CellChat package^33^. These regions were chosen to capture distinct regenerative contexts and thereby provide an integrated view of the cell–cell communication events shaping dystrophic regeneration.

Ligand–receptor analysis of region 1 containing the actively fusing myoblasts (Cluster 8) (Fig. 3A – highlighted by dashed box), revealed the activation of multiple signaling pathways (Extended Data Fig. 3A), including collagen, laminin, and fibronectin pathways. FAPs, fibroblasts, and the collagen expressing clusters 0 were the main sources of these pathways within the area (Extended Data Fig. 3B). Beyond cell-autonomous signaling, Cluster 8 (fibers undergoing myoblast fusion) showed a strong interaction with FAPs highlighting the crucial role of this cell type in providing ECM components needed for muscle regeneration (Extended Data Fig. 3B and Fig. 3B). These interactions in Cluster 8, which encompass committed and fusing myoblasts, were predominantly mediated by *CD44* (Extended Data Fig. 3C), which displayed a higher expression within the region when compared to SDC4 or DAG1 (Fig. 3C). This aligns with previous studies showing the role of CD44 in early myogenesis^34^ in contrast to SDC4, which primarily functions in MuSC activation^35^.

**Figure 3.**
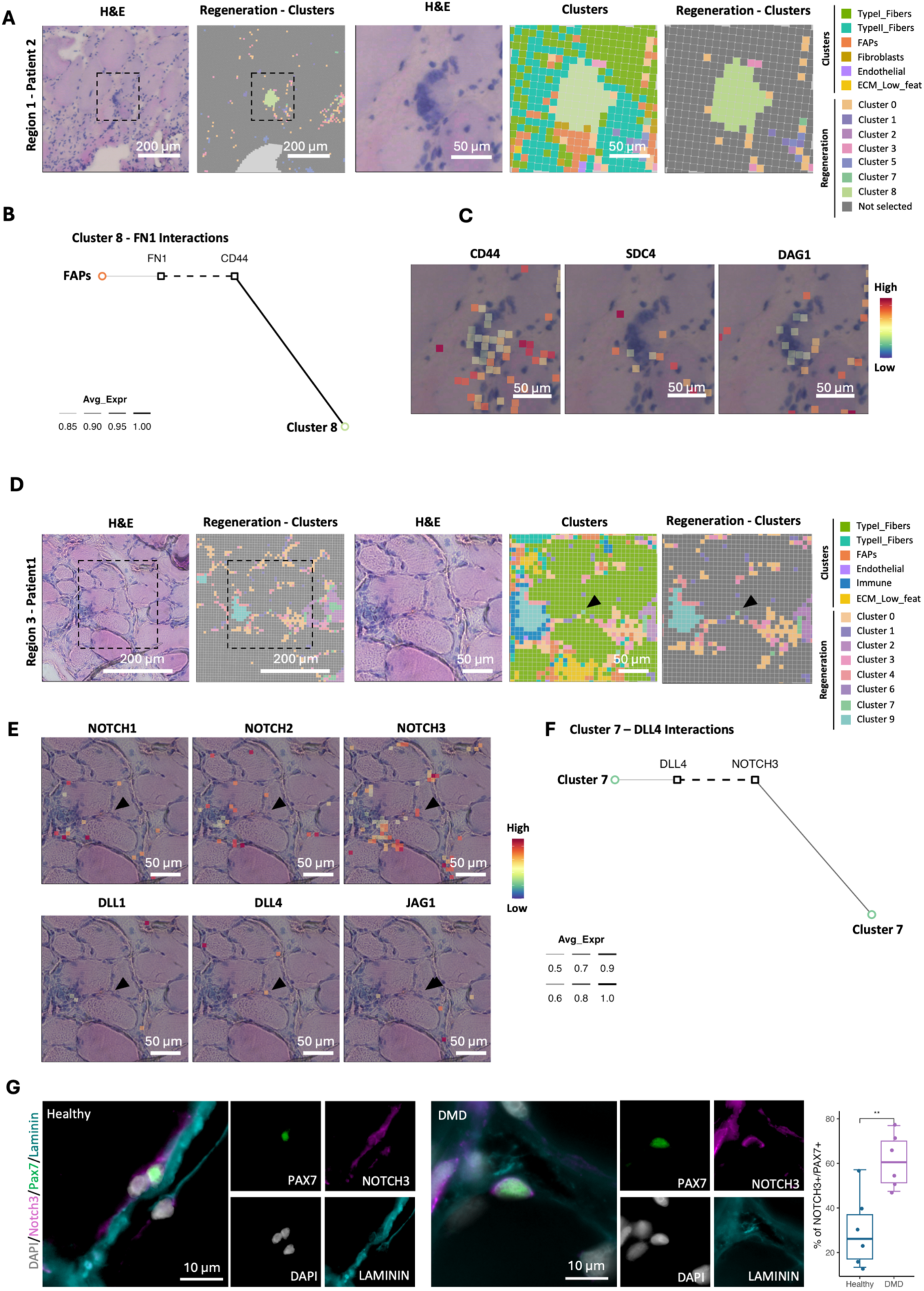
– Cell–cell communication analysis reveals stage-specific myogenic interactions and highlights NOTCH3 upregulation in activated satellite cells. A) Selected region 1 surrounding Cluster 8: H&E staining and spatial distribution of regenerative subclusters (left). Zoomed-in regions showing H&E staining and the combined spatial distribution of regeneration subclusters and cell type clusters (right). B) FN1-mediated interactions within the regenerative area around Cluster 8 cells highlighted in (A). C) Relative expression of CD44, SDC4, and DAG1 within the regenerative area in (A). D) Selected region surrounding Cluster 7: H&E staining and spatial distribution of regenerative subclusters (left). Zoomed-in regions showing H&E staining and the combined spatial distribution of regeneration subclusters and cell type clusters. E) Relative expression of NOTCH1, NOTCH2, NOTCH3 DLL1, DLL4 and JAG1 within the regenerative area in (D) Location of Cluster 7 on the image is marked by the arrow. F) DLL4-mediated interactions within the selected regenerative area around Cluster 7 cells in (D). Scale bars: 200 µm (overview images) and 50 µm (zoomed-in regions). G) Microscopy images of muscle sections from healthy individuals and DMD patients stained for Pax7, Notch3, Laminin, and Hoechst (left) scale bar 10 µm. Pax7–Notch3 colocalization was quantified as the number of Pax7⁺ cells also expressing Notch3 (Healthy n=6; DMD n=6, two-tailed unpaired t-test). Boxplots show the 75th, 50th and 25th percentiles; whiskers extend to values within 1.5 × the interquartile range, * p < 0.05.

Interestingly, we identified a region where Clusters 2 (regenerating fibers) was in close proximity to an area of immune infiltration (Cluster 9, marked by the expression of CD74; Region 2, Extended Data Fig. 4A-B). Analysis of incoming and outgoing signaling interactions within and among these clusters revealed a strong predominance of Cluster 9 as key signaling player (Extended Data Fig. 4C). Among the different active pathways, we detected *TWEAK* (*TNFSF12*) signaling (Extended Data Fig. 4D-E), linking the immune cluster to Cluster 2 (regenerating fibers). Several studies have suggested that TWEAK may negatively regulate myogenic differentiation^36,37^. Interestingly, its receptor, FN14 (*TNFRSF12A*), may have a TWEAK-independent role, as its deletion delays muscle regeneration^36^, and its silencing reduces myogenic regulatory factors^38^. In the selected region, *FN14* presented higher and more widespread expression than *TWEAK*, suggesting that, in specific areas, *FN14* might exert its function independently (Extended Data Fig. 4F). Among the different samples, overall *FN14* expression was highest in patient 1, where regenerating areas were most abundant, while it was lower in patient 2 (Extended Data Fig. 4G).

Finally, we examined a region surrounding Cluster 7 (enriched in activated satellite cells) to investigate active signaling pathways (Region 3; Fig. 3D). For Cluster 7, our analysis identified several active pathways including Collagen and Laminin, non-canonical WNT and NOTCH (Extended Data Fig. 5A). We focused on Notch signaling, which has been shown to play a key role in regulating myogenic progression and satellite cell quiescence in both healthy and diseased conditions^39–44,12,11,45–49^. CellChat analysis suggested the presence of potential autocrine signaling within Cluster 7 (Extended Data. Fig. 5B). When we examined the distribution of receptors and ligands within our region of interest, we detected higher levels of *NOTCH3* than of *NOTCH2* and *NOTCH1* (Fig. 3E) and DLL4 in Cluster 7. Intriguingly, Notch1 and Notch3 have been reported to be downregulated in the muscles of *mdx* mice^50^; however, in patient samples, *NOTCH3* is upregulated^12^. This discrepancy prompted us to further investigate the role of Notch signaling. Additional analysis of the imputed contributions of individual ligand‒receptor pairs revealed that the DLL4‒NOTCH3 was the most probable pair driving the potential signaling within the selected region (Extended Data. Fig. 5C and Fig. 3F). Next, we performed differential gene expression analysis. Our data revealed significant enrichment of *NOTCH3* in Cluster 7 compared with other regenerative clusters (Extended data Fig. 5D and Supplementary Table 3) as well as when compared to the other cell populations (Supplementary Table 3). In addition, *NOTCH3* was also upregulated in Cluster 4, endothelial cells, pericytes and smooth muscle cells (Supplementary Table 3). To validate these findings, we performed immunostaining across a set of human biopsies, confirming that NOTCH3 was more frequently expressed in satellite cells in Duchenne patient samples compared to healthy controls (Fig. 3G). To conclude, using our framework we charted the cellular communication networks in specific regenerating regions within dystrophic microenvironment. Our data highlight how FAPs and fibroblasts support early myoblast fusion through Collagen and Fibronectin signaling, and how TWEAK–FN14 interactions may further shape regenerative outcomes. Furthermore, our analysis prompted us to identify the upregulation of NOTCH3 in activated satellite cells and suggested NOTCH pathway as potentially deregulated within the dystrophic environment.

### NOTCH3 expression increases over time in MuSCs of the *mdx* mouse and its silencing improves myogenic differentiation

Since our human dataset provided only a static snapshot of regeneration, we generated a longitudinal multiomic dataset in the mdx mouse model to capture the dynamic changes in stem cell regulation and myogenic trajectory throughout disease progression. We performed paired snRNA-seq and snATAC-seq on the mdx dystrophic mouse model at different time points (2 weeks, 5 weeks, 6 months, and 14 months) (Fig. 4A). After quality control, the overall dataset contained 33,195 RNA and ATAC combined single-cell profiles (Fig. 4A). After clustering, the cells were annotated on the basis of canonical marker expression and assigned to major populations (Fig. 4B-C). In line with previous works in skeletal muscle ^8,51–56^, we identified cells representative of the different fiber types (1, 2A, 2B, 2X), components of the myotendinous junction, and neuromuscular junction. Within the mononuclear fraction, the major components were immune cells (myeloid cells and lymphocytes), MuSCs, FAP progenitors, endothelial cells, tenocytes, adipocytes, and smooth muscle cells and pericytes (SMMCs). Moreover, we were able to detect the whole regeneration continuum connecting satellite cells to the myofiber compartment, from myoblasts expressing *Megf10* and *Myod1* to *Myh3*-positive regenerative nuclei and *Flnc*-expressing regenerative fibers. The ATAC data were strongly correlated with the transcriptomic profile, suggesting that gene activity closely paralleled the RNA expression results (Fig. 4C). As we had identified NOTCH as a potentially deregulated signal in the human data, we aimed to determine whether *Notch3* upregulation was a direct consequence of dystrophin loss or rather a secondary effect of disease progression and impaired regeneration. To disentangle these possibilities, we next analyzed the expression of the Notch signaling pathway components in our mdx dataset. A comparison between actively regenerating muscle at 5 weeks and muscle at 6 months, when *mdx* muscles exhibits signs of impaired regeneration^57,58^, revealed a significant increase in *Notch3* expression in MuSCs (Fig. 4D). To further validate this finding, we performed qPCR on freshly isolated MuSCs (isolated as CD45^neg^ CD31^neg^ SCA1^neg^ ITGA7^pos^ VCAM1^pos^ – Extended Data Fig. 6A) and immunofluorescence staining of *tibialis anterior* sections, both of which confirmed increased *Notch3* expression in Pax7+ MuSCs at 6 and 14 months (Fig. 4E to G). Since previous studies have shown that *Notch3* downregulation in C2C12 and wild-type primary mouse myoblasts promotes myogenic differentiation^59^, we tested whether reducing *Notch3* levels in myoblasts from dystrophic 6-month-old mice could restore their differentiation potential. Primary myoblasts were transfected using siRNA against Notch3 or a scramble sequence while in growth condition with an approximate downregulation of 50% of Notch3 transcript (Extended Data Fig.6B). Cells were subsequently switched to differentiation media for 48 hours to induce myotubes formation. As predicted, silencing *Notch3* significantly improved myogenic differentiation, as evidenced by an increase in MyHC+ fiber area per mm^2^ (Fig. 4H-I) and the expression of the differentiation marker CKM (creatine kinase, muscle) (Fig. 4J). Altogether, these findings highlight a critical role for Notch3 upregulation in the progressive impairment of regenerative potential in dystrophic muscle.

**Figure 4.**
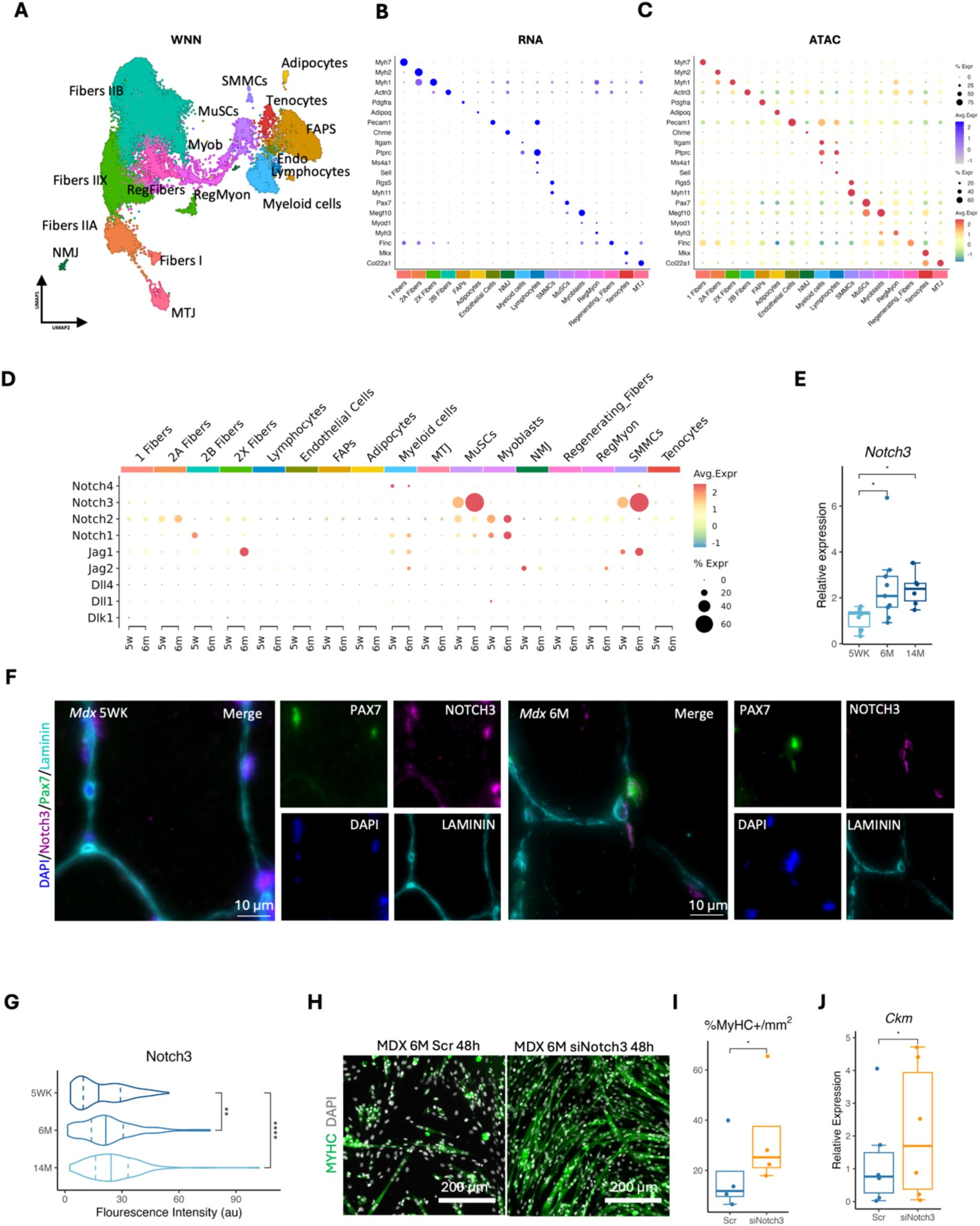
– Paired snRNA-seq and snATAC-seq on the mdx mice highlights Notch3 upregulation in MuSCs. A) Uniform manifold approximation and projection (UMAP) visualization and clustering of the weighted nearest neighbor (WNN) graph constructed by the integration of RNA and ATAC modalities; spots are color-coded on the basis of cluster assignment. Dot plots showing the RNA expression (B) and gene activity inferred based on chromatin opening (C) of the main representative genes for each cluster identified in the murine multiomic dataset. D) Dot plot showing the relative expression of Notch pathway ligands and receptors in the 5-month vs. the 6-month *mdx* dataset. E) Relative Notch3 mRNA expression in freshly isolated MuSCs from 5-week-old, 6-month-old, and 14-month-old *mdx* mice (5WK n = 10 6M n = 9, 14M = n = 6, Kruskal– Wallis/Dunn’s test). F) Microscopy images of TA muscle sections from 5-week-old and 6-month-old *mdx* mice stained for Pax7, Notch3, Laminin, and Hoechst. G) Notch3 fluorescence intensity in Pax7-positive Tibialis anterior from 5-week-, 6-month-, and 14-month-old *mdx* mice (n = 3 independent staining experiments, ≥25 Pax7⁺ cells quantified per condition in each experiment; data analyzed with a linear mixed-effects model followed by Tukey-adjusted pairwise comparisons). H) Microscopy image of 6-month-old dystrophic myoblasts stained for myosin heavy chain (MyHC) and Hoechst after Notch3 silencing and 48 h of differentiation scale bar 200 µm. I) Normalized MyHC-positive cell area after Notch3 silencing and 48 h of differentiation in 6-month-old dystrophic myoblasts (n = 4, two-tailed paired t-test). J) Relative *Ckm* mRNA expression of myoblasts from 6-month-old dystrophic mice after treatment with si*Notch3* or siSCR and 48 h of differentiation (n = 6, paired Wilcoxon test). Boxplots show the 75th, 50th and 25th percentiles; whiskers extend to values within 1.5 × the interquartile range. Violin plot show the 75th, 50th and 25th percentiles, The outline extends to the minimum and maximum data values. * p < 0.05, ** p < 0.01, *** p < 0.001.

**Figure 5.**
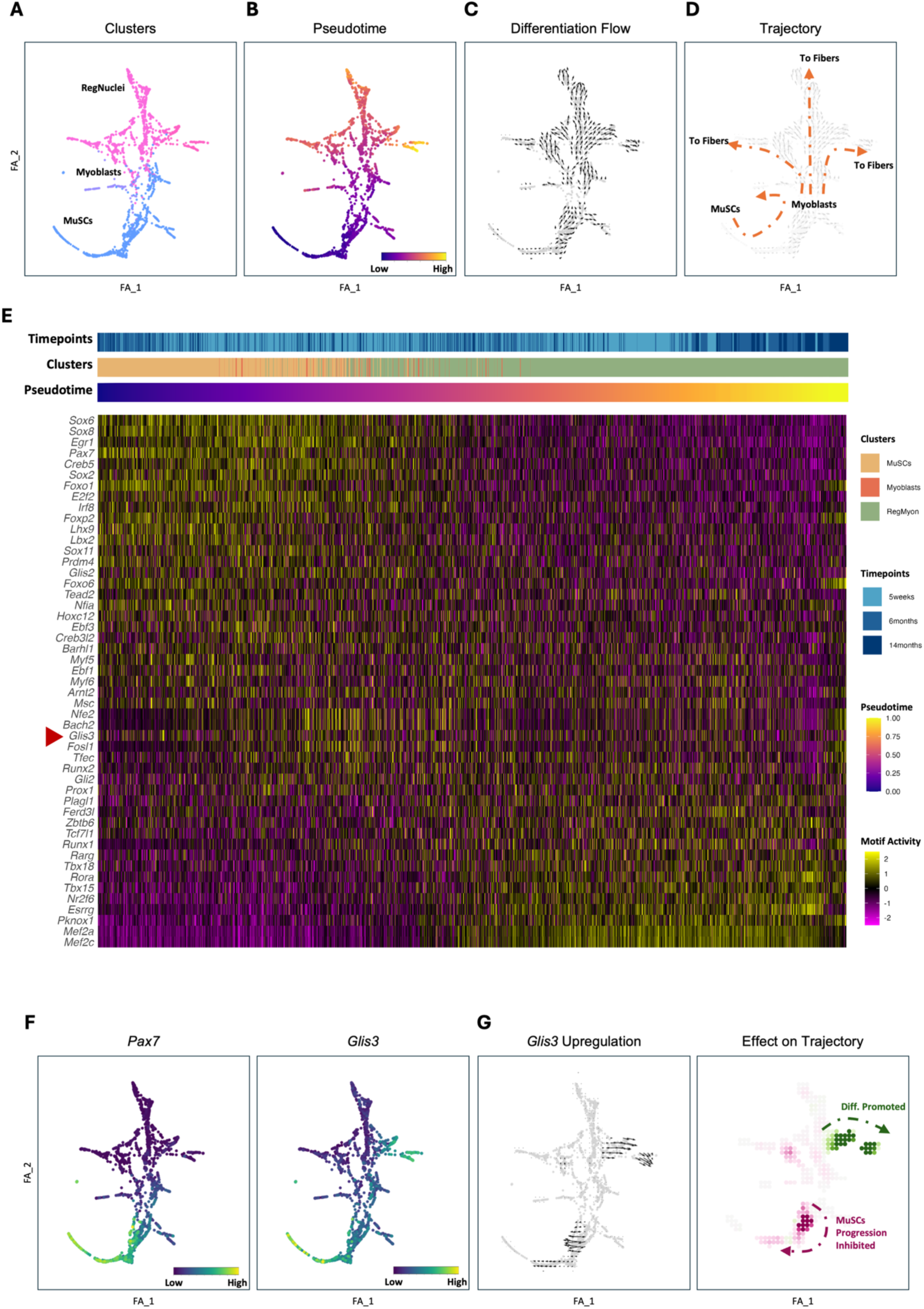
– CellOracle analysis identifies Glis3 as an intrinsic regulator of myogenic trajectory. A) Force atlas layout visualization of myogenic trajectory in *mdx*. The cells are color coded on the basis of their cluster (A) or their relative pseudotime value (B). The arrows in (C) describe the direction of differentiation flow. The diagram in (D) further illustrates the direction and endpoint of the differentiation process. E) Heatmap displaying imputed chromVar motif activity for transcription factors (TFs) identified by CellOracle, ordered by increasing correlation with pseudotime. The red arrow highlight *Glis3* F) Gene expression patterns of *Pax7* and *Glis3* within the myogenic trajectory. G) Changes in developmental flow upon *Glis3* upregulation (left). Schematic diagram further illustrating the effect on the myogenic trajectory.

**Figure 6.**
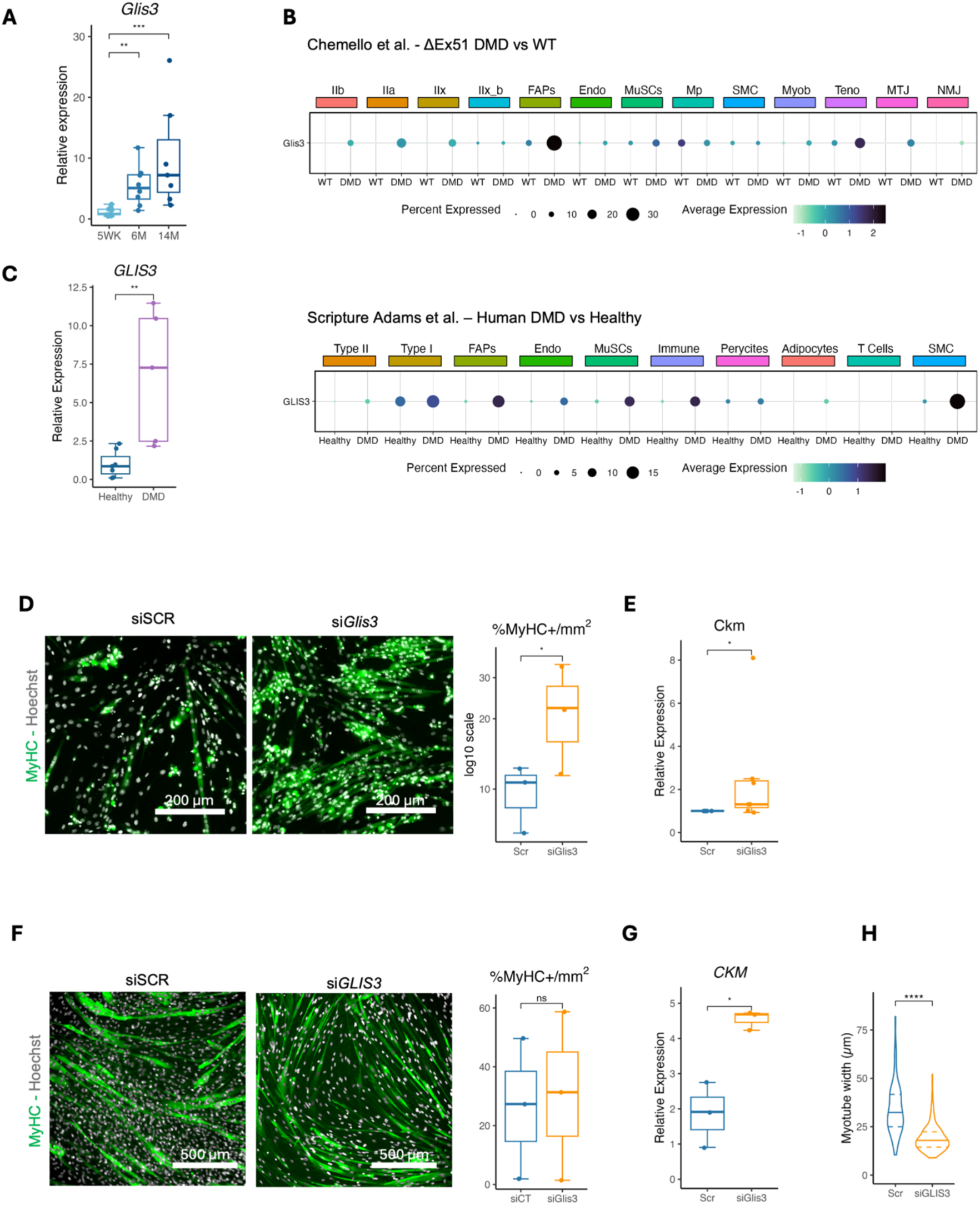
– *Glis3* expression in dystrophic muscle and functional impact on myogenesis. A) Relative *Glis3* mRNA expression in freshly isolated MuSCs from *mdx* mice at 5, 6, and 14 months (5WK n = 11, 6M n = 8, 14M n = 7, Kruskal–Wallis/Dunn’s test). B) Dot plot showing *Glis3* expression across different muscle populations in the ΔEx51 DMD mouse model (top) and DMD patient biopsies (bottom). C) Relative *GLIS3* mRNA expression in dystrophic and healthy human skeletal muscle samples (Healthy n = 7, DMD n = 5, two tailed Mann–Whitney U). D) Fluorescence microscopy image of MuSCs from 6-month-old dystrophic mice stained with MyHC and Hoechst after treatment with si*Glis3* or control siRNA (siSCR) and 48 h of differentiation scale bar:200 µm. The graphs represent the normalized MyHC-positive cell area (n = 3 two-tailed paired t-test)) (right). E) Relative *Ckm* mRNA expression of MuSCs from 6-month-old dystrophic mice after treatment with si*Glis3* or siSCR and 48 h of differentiation (n = 7, two-tailed paired t test). F) Representative images of patient-derived cell lines stained with MyHC and Hoechst after *GLIS3* silencing (or control treatment) and 4 days of differentiation scale bar:500 µm. The graphs represent the normalized MyHC-positive cell area (n = 3 independent cell lines; two-tailed paired t-test performed on log-transformed values) (right). G) CKM mRNA levels in siGlis3 and siSCR cells after 4 days in differentiation media (n= 3 biological replicates each point represents the average of two independent technical replicates, two-tailed paired t test). The values in G) are presented as relative increases compared with baseline (nontransfected samples) for each specific cell line. H) Myotube width quantification after 4 days of differentiation (n = 3 independent cell lines; ∼100 myotubes quantified per condition in total across the three lines; data analyzed using a linear mixed-effects model followed by Tukey-adjusted pairwise comparisons). Boxplots show the 75th, 50th and 25th percentiles; whiskers extend to values within 1.5 × the interquartile range. Violin plot show the 75th, 50th and 25th percentiles, The outline extends to the minimum and maximum data values. * p < 0.05, ** p < 0.01, *** p < 0.001.

### In silico Gene regulatory network inference highlights intrinsic regulators of the myogenic trajectory

ST analysis revealed key cellular interactions and signaling within the regenerating niche. To further explore intrinsic defects within the stem cell compartment leading to impaired regeneration, we next investigated the changes in transcription factors driving myogenic transitions. To achieve this goal, we used our single nucleus multiome dataset, and applied CellOracle^60^, a computational framework, which integrates epigenetic and transcriptomic data to predict how TF perturbations influence cell differentiation. We focused our analysis on MuSCs, myoblasts, and regenerative nuclei from the 5-week, 6-month, and 14-month time points. The cells were mapped onto a force-directed graph, ordered by pseudotime, and used to construct a developmental gradient vector field illustrating the trajectory of differentiation from MuSCs to regenerative nuclei (Fig. 5A to D). CellOracle analysis identified a set of TFs as potential regulators of myogenic progression (Supplementary Table 4). To determine the stage at which each TF was most active, we ranked them on the basis of the correlation between pseudotime values and their inferred motif activity (estimated using chromVar^61^) from low (high activity at low pseudotime values, i.e., early stages) to high (high activity at high pseudotime values, i.e., late stages). As expected, our analysis highlighted several well-known myogenic TFs, including *Pax7* and *Myf5* for early stages and *Mef2c* and *Mef2a* for later differentiation stages (Fig. 5E). To assess how disease progression impacts regulators of the myogenic trajectory, we focused on TFs that were differentially expressed at 6 months compared with 5 weeks (Supplementary Table 5). Among them, we focused on GLIS family zinc finger 3 (Glis3), a nuclear protein with five C2H2-type zinc fingers^62^, as it showed the highest upregulation among the significantly altered genes (p-value <0.05). While Glis3 functions as both a transcriptional activator and repressor and plays a regulatory role in various cells and tissues—including pancreatic beta cells, the thyroid, the liver, and the kidney^63,64^ — its role in skeletal muscle remains largely unexplored. Notably, however, its Drosophila ortholog *lame duck* (also known as *gleeful*) is involved in the specification and function of fusion-competent myoblasts^65,66^, thus suggesting a potential link with myogenesis. In our dataset, *Glis3* closely overlapped *Pax7* expression, suggesting a possible role in the early stages of differentiation from MuSCs to regenerative nuclei. Additionally, *Glis3* was enriched in regenerative nuclei, aligning with a potential function in myoblast fusion (Fig. 5F). Interestingly, Glis3 expression increased during later stages of disease, not only in MuSCs but also across the entire myogenic continuum (Extended Data. Fig. 7A). This observation raised the question of whether *Glis3* upregulation actively contributes to regeneration impairment. To assess the potential functional impact of increased *Glis3* expression, we performed in silico TF upregulation analysis. Simulation of a twofold increase in *Glis3* expression predicted a delay in the transition from MuSCs to myoblasts coupled with an accelerated differentiation of regenerative nuclei (Fig. 5G). Together, these results suggest that Glis3 upregulation may shorten the amplification phase of committed myoblasts, ultimately reducing the pool of differentiating cells.

### Glis3 deregulation in dystrophic muscle interferes with the myogenic trajectory

To further investigate the role of Glis3, we first aimed to confirm its upregulation in dystrophic MuSCs during disease progression. In line with the snRNA-seq data, quantitative PCR analysis of freshly FACS-isolated cells revealed a significant increase in *Glis3* expression in satellite cells isolated from 6-month and 14-month-old mice compared to those isolated from 5-week-old mice (Fig. 6A). We also analyzed publicly available snRNA-seq datasets from an additional dystrophic mouse model^8^ (ΔEx51 DMD) and human DMD patients^14^ and determined that *Glis3* was upregulated not only in MuSCs but also in FAPs and tenocytes, with an even more generalized trend in human snRNA-seq data (Fig. 6B). We validated this observation through qPCR on bulk RNA from dystrophic and healthy human muscle samples (Fig. 6C).

Once the expression pattern of Glis3 in the dystrophic muscle was confirmed, we next sought to validate our in-silico prediction of its impact on the MuSC trajectory. To achieve this goal, we isolated satellite cells from 6-month-old *mdx* mice, cultured and transfected them with an siRNA against *Glis3* or a nontargeting sequence. To rule out potential off-target effects on related sequences, we also assessed the relative expression levels of other Glis and Gli family members. At 48hrs Following siRNA transfection, *Glis3* transcript showed approximately a 50% reduction while none of the tested off-target transcripts showed significant changes, confirming the specificity of the siRNA (Extended Data Fig. 7B-C). Upon differentiation, we measured the relative percentage of MyHC+ area, and the levels of the *Ckm* transcript (Fig. 6D-E). These parameters exhibited increases upon *Glis3* downregulation, supporting the idea that a downregulation of Glis3 could lead to an overall improved differentiation.

Finally, we sought to understand whether the role of *Glis3* was conserved in humans. To do so, we silenced GLIS3 in muscle cells derived from DMD patients. As expected, due to the inherent variability in cellular composition from muscle explants, these cell lines showed significant differences in myogenic potential (Fig. 6D and Extended Data Fig. 8A). We transfected the cells in growth medium and, 48 hours later, we shifted them to differentiation media for an additional four days. Efficient knockdown was confirmed by qPCR, showing approximately a 65% reduction in GLIS3 transcript levels (Extended Data Fig. 8B). MyHC-positive area showed an increasing trend upon GLIS3 depletion, although it did not reach statistical significance, and CKM expression was significantly increased (Fig. 6F and G). Moreover, myotubes from the si*GLIS3* group displayed a decreased relative width compared with those of their control counterparts (Fig. 6H; data for individual cell lines are presented in Extended Data Fig. 8C), in line with an additional role for GLIS3 in maturation. Taken together, these data indicate that GLIS3 expression is elevated in dystrophic muscle and specifically in MuSCs, where its upregulation interferes with myogenic progression. Importantly, GLIS3 appears to act at multiple points in the differentiation program, not only during early activation but also throughout myotube maturation.

## Discussion

Here, we generated a new resource to study impaired muscle regeneration in dystrophic muscle, leveraging spatial and multiomic single-nuclei-data. We identified the upregulation of NOTCH3 and GLIS3 as barriers to the correct execution of the myogenic program. Our analysis revealed increased NOTCH3 expression in MuSCs from patients and from late stage mdx mice and demonstrated that its silencing *in vitro* enhances myogenic differentiation. Furthermore, using a combination of *in-silico* TF perturbation and data from human and murine samples we have identified that GLIS3 is a potential regulator of myogenic trajectory and that its upregulation in the dystrophic MuSCs leads to defective myogenic progression.

Using high-resolution spatial transcriptomics, we annotated distinct morphological and functional regions of skeletal muscle, identified the full regenerative continuum, and characterized the interactions between regenerative populations and their surrounding microenvironment. Through our analysis, we showed that multiple signaling pathways, such as those involving Laminin, Collagen, and Fibronectin, are mediated through CD44 rather than SDC4 in myofibers undergoing nuclear fusion. This finding is in line with the known role of CD44 in chemotaxis and early myogenesis^34^ and the fact that *SDC4* is involved primarily in MuSC activation and amplification^35^. Furthermore, these interactions were predominantly mediated by FAPs, consistent with previous reports providing additional validation for our analysis^15^. In addition, we detected the activation of the TWEAK signaling pathway, which links fibers undergoing maturation and the immune compartment. Several previous *in vitro* and *in vivo* studies have suggested *TWEAK* as a negative regulator of myogenic differentiation^36,37^. Additionally, this signaling pathway is upregulated in inclusion body myositis^37^, and its blockade improved muscle function and histology in a murine model of myotonic dystrophy type 1^67,68^. Interestingly, its receptor, FN14 (*TNFRSF12A*), may have a TWEAK-independent role, as its deletion delays muscle regeneration^36^, and its silencing reduces myogenic regulatory factors^38^. In our dataset, *FN14* showed higher and broader expression than TWEAK, indicating potential independent functions. A reanalysis of the Affymetrix dataset of Duchenne patients from Pescatori et al.^69^ presented by Burkly and colleagues^70^ identified *FN14* as upregulated in quadriceps muscles from young DMD patients, linking it to an environment where regenerative potential is not yet completely exhausted. Similarly, in our dataset, *TNFRSF12A* expression was highest in Patient 1, where regenerating areas were the most abundant. Our results confirm previously published findings, demonstrate that HD spatial transcriptomics can be effectively used to identify key signaling pathways, thereby advancing our understanding of DMD pathology and aiding the search for potential therapeutic strategies.

Several studies have shown the role of NOTCH signaling as a regulator of satellite cell quiescence and myogenic progression and in skeletal muscle under both healthy^39–44,12,11,45–49^ and diseased conditions. In our analysis, NOTCH3-mediated signaling emerged as a key pathway in activated dystrophic satellite cells. We detected NOTCH3 upregulation in MuSCs from patients and from mdx mice aged 6 months and older. Although this increase may appear to contrast with what previously reported^50^, the discrepancy likely arises from differences in the cell types analyzed. Our study focused on freshly isolated MuSCs, whereas prior studies assessed cultured primary myoblasts.

Collectively, our data suggest that Notch3 might reach levels able to interfere with the correct execution of the myogenic program in MuSCs only at an advanced stage of the disease and that this effect is at least partially reversible as it can be mitigated by silencing its transcript.

This is in line with the literature, where it has been shown that Notch3 overexpression inhibits satellite cell differentiation, while its silencing improves myogenic differentiation in C2C12 cells^59^. This model is further supported by the late onset of hyperplasia in *mdx/Notch3*^-/-^ mice ^43^. In these mice, although the early regenerative wave characteristic of the mdx model occurs normally, the absence of Notch3 does not lead to noticeable differences in muscle histology at 8 weeks^43^. This indicates that Notch3 does not significantly influence muscle regeneration at this stage. On the other hand, as the *mdx/Notch3*^-/-^ mice age, muscle size strikingly increases despite the limited regeneration typically observed in adult mdx^43^. Strongly suggesting an involvement of Notch3 at an advanced stage of the disease. Our findings, along with these previous observations, indicate that attenuating Notch3-mediated signaling at late stages of the disease may sustain regenerative capacity in dystrophic muscle, maintaining tissue integrity and potentially extending the therapeutic window for effective gene therapy. However, for Notch3 to be a viable target, it will be essential to further investigate the potential impact on endothelial cells and smooth muscle cells, which also express Notch3 and could be negatively affected, especially by long-term treatments.

While spatial transcriptomics provided a broad view of the different interactions within regenerative regions, we also aimed to identify intrinsic transcriptional changes within MuSCs that drive disease progression. To achieve this goal, we leveraged our multiomic dataset to trace the myogenic trajectory and identify TFs that may contribute to repair impairment within DMD muscle. Among the candidates identified, we focused on Glis3, a member of the Gli-like family of Krüppel-like zinc finger TFs. Glis3 has been primarily studied in the context of pancreatic β-cell development, kidney function, diabetes, and testis development^64^. In our dataset, Glis3 expression was elevated in MuSCs and partially overlapping with Pax7. Intriguingly, this pattern is also observed in the testis where both have been reported to be expressed in spermatogonial stem cells ^63,71,72^. Although its role in skeletal muscle remains largely unexplored, two related genes, Glis1 (GLIS Family Zinc Finger 1) and Gli3 (GLI Family Zinc Finger 3), have been recently reported to play a role in MuSC fate and self-renewal. The first one, *Glis1*, which shares 93% homology in the Zinc Finger Domain (ZFD)^62^, has a pro-adipogenic role and promotes MuSCs differentiation into brown adipocytes^73^. The other one, *Gli3*, which shares less than 70% homology in the ZFD^62^, has been implicated in MuSCs entry into G-Alert and self-renewal^74^. Notably, *Gli3* deletion enhances hypertrophy and increases satellite cell numbers upon single and repeated injury^74^. Our data suggest that, despite the homology, *Glis3* exerts a distinct function in dystrophic MuSCs as its silencing leads to a different phenotype: enhanced progression to the myoblast stage but dampened myotube diameter, pointing to an additional role in myotube maturation. This is in line with its upregulation following strength training^75^ and the implication of its Drosophila ortholog, gleeful, in the fusion of competent myoblasts^66^. Taken together, our findings suggest that Glis3 is a previously overlooked regulator of MuSCs during skeletal muscle regeneration, with significant implications for dystrophic disease progression. Future studies should focus on its downstream effectors to determine whether targeting GLIS3 could preserve myogenic potential while minimizing effects on maturation.

In conclusion, this study provides a high-resolution spatial view of the regenerative landscape in dystrophic muscle, offering new mechanistic insights into the cellular and molecular alterations that underlie repair impairment in DMD. By investigating the full regenerative continuum and analyzing the changes in different TFs regulating myogenic progression, we identified NOTCH3 and GLIS3 as contributors to the progressive decline in muscle regeneration. Importantly, modulating these pathways enhanced differentiation, highlighting their potential as therapeutic targets. Altogether, our results lay the groundwork for future development of novel therapeutic strategies aimed at improving regeneration in dystrophic muscle. More broadly, our work illustrates how spatial and multiomic approaches can bridge the gap between cellular identity, tissue architecture, and disease progression, providing a powerful framework to investigate pathogenic mechanisms across a wide range of tissues and diseases.

## Limitations of the study

While ST offered us the opportunity to investigate local dynamics in the regenerative areas at high resolution, it should be considered that the relative sample size in our dataset (2 healthy donors and 2 DMD patients) limits the extrapolation of the spatial gene expression patterns observed and reduces statistical power for detecting subtle regional differences. It should also be noted that since our analysis relies on 8-µm bins for annotation and cell–cell interaction, we cannot resolve features below this resolution; as a result, bins may in principle capture signals from multiple adjacent cells. Lastly, the quality of DMD patient muscle samples varied, potentially introducing regional loss of specific transcripts due to mRNA degradation. While HD ST provides unprecedented spatial resolution, it is important to acknowledge that, like any emerging technology, it has inherent technical variability, including sensitivity to tissue processing artifacts. Together, these limitations underscore the need for future studies with larger, more diverse cohorts and standardized protocols to validate and extend our findings.

## Materials and Methods

### Muscle biopsy and ethical clearance

Human skeletal muscles were obtained from the Myobank-AFM, Paris, France (https://www.institut-myologie.org/en/recherche-2/myobank-afm/). Samples were obtained with patient or parental written consent, collected from surgical residue from patients undergoing surgery under the authorization of the Minister of Research, approval number AC-2019-3502, in accordance with French and European laws.

### Mouse models

Mice were handled in accordance with European Community guidelines. Experimental animal protocols were performed in accordance with the guidelines of the French Veterinary Department and approved by the Sorbonne Université Ethical Committee for animal experimentation (APAFIS #36186-2022032511143182 v4). In this study, 2 weeks, 5 weeks, 6 months and 14 months C57BL/10-*mdx*^76^mice were used.

### Cell lines and culture

Primary mouse myoblasts were cultured in growth medium Dulbecco’s Modified Eagle Medium (DMEM, Thermo Fisher Scientific, Waltham, MA, USA) supplemented with 20% fetal bovine serum (FBS, Thermo Fisher Scientific), 5 ng/ml basic fibroblast growth factor (Thermo Fisher Scientific) and 1% penicillin/streptomycin (Thermo Fisher Scientific) on Matrigel (1:100, Corning, NY, USA) coated plates under a humidified 5% CO2 atmosphere at 37°C.

Non-immortalized primary DMD patient derived cell lines 1,2, and 3 were obtained from the MyoLine Platform (Myology Institute, Paris, France). Cells were culture in Dulbecco’s Modified Eagle Medium (DMEM supplement high glucose and GlutaMAX, Thermo Fisher Scientific) supplemented with 16% medium 199 (supplement GlutaMAX, Thermo Fisher Scientific), 20% FBS (Thermo Fisher Scientific), 50ng/ml Gentamycin (Thermo Fisher Scientific), 25 µg/ml Fetuin (Thermo Fisher Scientific), 5ng/ml human Epidermal Growth Factor (Thermo Fisher Scientific), 0,5 ng/ml basic fibroblast growth factor (Thermo Fisher Scientific), 5 µg/ml Insulin (Merck) and 0,2 µg/ml dexamethasone (Merck) on Matrigel (1:100, Corning, NY, USA) coated plates under a humidified 5% CO2 atmosphere at 37°C.

### Primary myoblast transfection and differentiation

Reverse transfection was performed on primary myoblasts using Silencer Select siRNAs and RNAiMAX kit (both Thermo Fisher Scientific). For mouse experiments, were used Silencer Select siRNAs against Glis3 (s105279) and Notch3 (s70709). To minimize variability, silencing experiments for Glis3 and Notch3 were performed simultaneously using the same siCTR controls, ensuring consistent experimental conditions. The transfection complexes were prepared inside Matrigel-coated wells, after which the cells were seeded at density of 25 000 cells/cm2. Transfection complexes were incubated for 48 hours before switching to differentiation medium (DMEM with 5 % horse serum –HS, Eurobio Scientific, 1% penicillin/streptomycin). The cells were kept in differentiation medium for 24 – 48 –hours. For human experiments, Silencer Select siRNAs against GLIS3 (s46764) were used. The transfection complexes were prepared inside Matrigel-coated wells, after which the cells were seeded at density of 25 000 cells/cm2. Transfection complexes were incubated for 48 hours before switching to differentiation medium (DMEM with 5 % horse serum (HS), 10 µg/ml insulin and 50 µg/ml gentamycin). The cells were cultured in differentiation medium for 4 days.

### Satellite cell isolation and Fluorescence-activated Cell Sorting (FACS)

Primary Satellite cells were isolated based on the protocol from Liu and colleagues ^77^ with minor modifications. Briefly, hindlimb muscles from euthanized mice were dissected and minced, then incubated for 1 h at 37 °C with agitation in wash medium (Ham’s F-10 medium – Thermo Fisher Scientific, supplemented with 10% horse serum and 1% penicillin/streptomycin) containing collagenase type II (800 U/ml; Worthington Biochemical, Lakewood, NJ, USA). After incubation, samples were centrifuged (500 × g, 5 min), resuspended in wash medium supplemented with collagenase type II (1,000 U/ml) and dispase (11 U/ml; Roche, Basel, Switzerland), and incubated for an additional 30 min at 37 °C with agitation.

Following digestion, samples were filtered, centrifuged, and treated with red blood cell lysis buffer (Promega, Madison, WI, USA) according to the manufacturer’s protocol. Cells were then resuspended in wash medium and incubated with primary antibodies (see Key Resource Table) for 45 min. Prior to FACS, samples were passed through a 50 µm filter.

Labeled cells were sorted using a BD Aria cell sorter (70 µm nozzle), and muscle stem cells (MuSCs) were collected as CD31⁻ CD45⁻(Lin^-^) Sca1⁻ ITGA7⁺ VCAM1⁺. The resulting cell suspension was centrifuged (500 × g, 5 min, 4 °C) and either plated in growth medium or processed for RNA extraction.

### Immunostaining

Cells on CellCarrier Ultra microplates (Thermo Fisher Scientific) were fixed for 10 minutes in 4% paraformaldehyde (PFA, Electron Microscopy Sciences, Hatfield, PA, USA) and permeabilized for 10 minutes in 0.1% Triton X-100 (Euromedex, Tuileries, France). Samples were blocked for 30 minutes in blocking buffer (1% BSA –Eurobio Scientific-in 1xPBS) at room temperature (RT). Primary antibodies were diluted in blocking buffer and incubated at RT for one hour (Table 1). The samples were washed three times with PBS and then incubated with secondary antibodies (Table 1) and Hoechst diluted in PBS for 30 min. Mice TA and EDL muscles were frozen in isopentane (Avantor, PA, USA) cooled with liquid nitrogen. Frozen sections of 10 μm thickness were fixed in 4% PFA for 10 minutes and permeabilized in pre-chilled methanol for 6 minutes at –20 °C. For anti-Pax7 staining, antigen retrieval was performed by heating slides at 95 °C in Antigen Unmasking Solution (Vector Laboratories, Newark, CA, USA) for 15 minutes. After antigen retrieval, samples were cooled on ice and washed once with PBS. Samples were then blocked for 2 hours in blocking buffer (5% BSA, 10% horse serum, 0.1% Triton X-100) at RT. Primary antibodies (Table 1) were diluted in blocking buffer and incubated overnight at +4 °C. The next day, the samples were washed three times with PBS and incubated with secondary antibodies (Table 1) for 1 hour followed by Hoechst staining for 10 minutes. After washing three times with PBS, a coverslip (Knittel Glass, Braunschweig, Germany) were mounted on microscopy slides (J1800AMNZ, Epredia) using mounting medium (Thermo Fisher Scientific).

### Microscopy and image analysis

Microscopy was performed using an inverted Nikon Eclipse Ti2 microscope controlled by NIS-Elements software (Nikon). Images were acquired with ORCA Flash4 sCMOS camera and with 40x oil-immersion or 20x air objective lenses. Confocal images were acquired using an inverted Nikon Ti2 eclipse microscope equipped with a motorized stage and a Yokogawa CSU-W1 (Tokyo, Japan) spinning disk head controlled by Metamorph software (Molecular Devices, San Jose, CA, USA). Images were acquired with a Prime 95 sCMOS camera (Photometrics, Tucson, AZ, USA) and with a 100x or 40x oil-immersion objective lenses. Myotube area and fluorescent intensity were analyzed with ImageJ Fiji software (Schindelin et al., 2012).

### Quantitative Reverse Transcription PCR

RNA was isolated using Trizol (Thermo Fisher Scientific) and Direct-zol RNA MicroPrep (Zymo Research, Irvine, CA, USA) according to the manufacturer’s instructions. The cDNA was synthesized using QuantiTect Reverse Transcription kit (Qiagen, Hilden, Germany). Quantitative real-time PCR was carried out using the FastStart Universal SYBR Green Master (Roche). Each sample was amplified in duplicate in a thermal cycler (QuantStudio 7 Pro, Thermo Fisher Scientific) and transcript levels were normalized to ribosomal subunit protein (Rplp) or Glycéraldéhyde-3-phosphate dehydrogenase (GAPDH) for human samples. Relative mRNA levels were calculated using the ΔΔCT method. The primer sequences are listed in the Key Resource Table.

### HD Spatial Transcriptomics

Fresh Frozen muscle human samples were cryo-sectioned at 10 μm thickness at –22°C. For each sample, a single cross-section was placed on a superfrost slide. The slide was transferred onto dry ice and brought to room temperature (RT) for 5 minutes, followed by fixation in cold methanol for 30 minutes at –20°C. Subsequently, the slide was covered with isopropanol at RT for 1 minute, after which hematoxylin and eosin (H&E) staining was performed according to the manufacturer’s protocol. After image acquisition, the slide was destained and mounted on the 10X support. Muscle tissue was then subjected to probe hybridization and ligation. Probes were subsequently released and captured by the Visium HD slide using the CytAssist instrument (10x Genomics). Following probe extension, cDNA was amplified to generate a spatial gene expression library. Sequencing was performed on an Illumina Nextseq 2000. Loupe Browser v8 (10x Genomics) was used to align high resolution images to Cytassist-generated image.

### HD spatial transcriptomics data analysis

Raw sequence data were processed using Space Ranger (3.0.0) and aligned to the human reference genome (GRCh38-2024-A). Downstream analysis of the dataset was performed using Seurat v5^19^. Data matrix and aligned images were loaded into a Seurat object using the command Load10X_Spatial with bin.size parameter set to 8. A high-resolution image was then loaded using the command Read10X_Image and added to Seurat object. A sketch assay was created (ncells = 5000, method = “LeverageScore”) to perform dataset integration using RPCA method with the following parameters: dims = 1:30, k.anchor = 20, reference = which(Layers(HD_merged, search = “data”) %in% c(“data.CTR3”) (healthy control 1). Initial dataset clustering was performed using FindClusters command with a resolution parameter of 0.5. Cluster were then transferred from the sketched assay to the full Spatial.008um dataset using ProjectData() to define the main populations within the integrated dataset. Each cluster was then analyzed separately and re-clustered. Subclusters were assigned to main populations based on canonical markers. Cell–cell communication and ligand–receptor interaction analyses were performed using the R package CellChat^33^. Briefly, the CellChat object was created from a subset of the integrated seurat object containing only the selected region of interest. Spatial information was loaded into CellChat object as described in CellChat vignette. Communication probability was calculated using computeCommunProb function with the following parameters type = “truncatedMean”, population.size = FALSE, distance.use = TRUE, contact.range = 10, trim = 0.1,k.min = 5, scale.distance = 0.12, contact.dependent = TRUE.

### Nuclei extraction, sorting and library preparation for 10x chromium single cell multiome ATAC and gene expression

Hindlimb muscles from two mice per age group were pooled for combined snRNA and snATAC-seq with the 5-week and 6-month groups also including an additional technical replicate. Nuclei were extracted based on the protocol described in^54^ with minor modifications. The NP40 Lysis buffer, 0.1 Lysis Buffer and wash buffer were prepared according to manufacturer’s instructions (Nuclei Isolation from Complex Tissues for Single Cell Multiome ATAC + Gene Expression Sequencing – CG000375 10x Genomics). Briefly, muscles were minced and placed for 5 minutes into 2 mL of NP40 lysis buffer. During incubation tissue was cut into smaller pieces, with occasional agitation of the solution. At the end of incubation 2 mL of wash buffer was added. The solution was transferred into a 7 mL dounce and tissue was subjected to 25 strokes with piston A. The solution was then transferred to a 5 mL Eppendorf tube. The sample was filtered through a 100 µm filter, followed by a 40 µm filter, into a 50 mL Falcon tube. Next samples were centrifuged at 500g for 10 minutes at 4°C on a refrigerated centrifuge (acceleration of 8 and a deceleration of 3). The supernatant was discarded, and 1 mL of wash buffer v3 (1%BSA, 1 U/µL RNAse inhibitors in PBS) was added. The solution was incubated on ice for 5 minutes. The pellet was resuspended, and the sample was centrifuged again under the same conditions. The supernatant was discarded, and the pellet was resuspended in 500 µL of wash buffer, to which 10 µL of 7-AAD was added. The solution was incubated for 5 minutes on ice before being filtered through a 20 µm filter into a FACS tube. A FACSAria sorter was used to sort 800,000 7-AAD-positive nuclei with a 100 µm nozzle at a cold block setting. In the collection tube (15 mL), were added 500 µL of 10% BSA and 5000 U of RNase Inhibitors (Protector –Roche). A FACSAria sorter was used to sort 800,000 7-AAD-positive nuclei with a 100 µm nozzle at a cold block setting. After sorting, the collection tube was adjusted to 5 mL with 1X PBS. For nuclear membrane permeabilization, the collected sample was centrifuged at 500g for 7 minutes at 4°C (acceleration = 8, deceleration = 1), ensuring the tube was properly balanced. The supernatant was discarded carefully to avoid disturbing the pellet. The pellet was resuspended in 100 µL of 0.1X Lysis Buffer, up and down five times, and incubated on ice for 2 minutes. Next, 1 mL of wash buffer was added. The sample was centrifuged again under the same conditions for 7 minutes. The supernatant was discarded, and the pellet was resuspended in 50 µL of Diluted Nuclei Buffer and kept on ice. The final concentration of nuclei was measured using the Nucleocounter NC200 ChemoMetec (Allerod, Denmark). The required volume for loading 12,500 nuclei was calculated, and the nuclei count was confirmed using a Malassez hemocytometer. Next, nuclei were loaded into GEM Chip J with the Chromium Next GEM Single Cell Multiome Reagent Kit to generate GEMs with the Chromium Controller (10x Genomics). All subsequent GEM incubation post-GEM cleanup, pre-amplification, ATAC library construction, cDNA amplification and GEX library construction steps were performed according to the manufacturer’s instructions. Libraries were sequenced according to manufacturer’s instructions on a NextSeq 500 instrument.

### Single-nuclei RNA and ATAC sequencing data analysis

Raw sequence data for mouse samples were processed using Cell Ranger ARC (2.0.0) and aligned to the mouse reference genome (mm10). Further analysis of the dataset was performed using Seurat 5 ^19^. Each sample was preprocessed with SoupX package and the modified matrices were used to generate Seurat objects. Normalization, scaling, and quality control were performed separately on each object before integration. Nuclei with fewer than 350 detected genes or with >20% mitochondrial gene content were removed from the dataset. Different datasets were integrated with PrepSCTIntegration() function and using 5000 features for integration. The integrated dataset was normalized using SCT. After the first integration, snATAC-seq data were integrated with snRNA-seq data. Unsupervised clustering identified 28 clusters in the combined snRNA/ATAC-seq. Clusters were then manually annotated into different cell types based on canonical markers expression. Uniform manifold approximation and projection (UMAP) was applied to visualize the individual clusters in the dataset. After clustering and cell population annotation, differentially expressed genes between age groups were identified using the FindAllMarkers() function with the following parameters: only.pos = TRUE, min.pct = 0.25 and logfc.threshold = 0.25. Cell oracle analysis was performed using default parameters. Glis3 upregulation has been set to 0.8 (equivalent to a 2-fold induction).

## Statistical analysis

Statistical analyses were performed using R. The exact sample sizes, number of measurements and statistic test used for each experiment are indicated in the figure legends. Datasets following a normal distribution were analyzed with two-tailed paired Student’s t-test or grouped two-way ANOVA followed by Tukey’s post hoc test. Non-parametric datasets were analyzed with Wilcoxon test (paired samples), Mann–Whitney U test (unpaired samples) or Kruskal–Wallis test followed by Dunn’s test for multiple comparisons. Datasets requiring aggregation of repeated measures for each biological replicate were analyzed using a mixed linear model. For figure 4G age was included as a fixed effect and experiment as a random intercept, and for figure 6H condition was modeled as fixed effect and cell line as random intercept. P-values ≤ 0.05 were considered statistically significant and are marked with symbols in the figures.

## Figure Design

Figure 1A was created with BioRender (https://biorender.com/) licensed to LG

## Acknowledgments

We thank the patients and their families for consenting to the use of donated tissue samples for this research. In addition, we would like to thank the MyoBank and MyoLine platforms for collecting and kindly providing patient-derived samples and myoblasts. We would like to thank Dr. Pietri-Rouxel and Dr. Gentil for their kind help and support in the initial phase of the project. We thank the CyPS Facility (Sorbonne University, Paris) for their technical support. We would also like to acknowledge the Genomi’C and CyBio platforms (Cochin Institute, Paris) for generating the multiomic dataset and the Histomics and Igenseq platforms (Paris Brain Institute, Paris) for generating the spatial dataset. This work was supported by the ANR (Agence Nationale de la Recherche) grant ANR-20-CE14-0048. LV has been supported by the Instrumentarium Science Foundation and by ANR-20-CE14-0048. FCG was financed through support of the France Relance National Program and the Marie Curie Postdoctoral Fellowship (GAP: 101109098).

## List of supplementary tables

Supplementary Table 1 – List of dystrophic muscles obtained from patients undergoing surgical operations used for Spatial transcriptomics, Pax7+/Notch3+ count and GLIS3 whole muscle RNA levels.

Supplementary Table 2 – Percentage of clusters per dataset

Supplementary Table 3 – Lists of differentially expressed genes identified within regenerating clusters only, and across the entire dataset including all populations (regeneration and non-regeneration clusters)

Supplementary Table 4 – List of TFs identified by CellOracle

Supplementary Table 5 – List of TFs differentially expressed at 6 months and 5 weeks

## Contributions

L.G. supervised the study. L.G. and L.V. designed the experiments and analyzed the results. C.D., L.S., F.G., and C.P. performed experiments and analyzed data. H.B. and L.V. conducted the FACS experiments. K.V., B.C., I.B., and S.B. carried out the multiomic experiments under the supervision of A.M., L.F., and M.C. S.V. collected and stored the muscle samples. K.M. isolated the patient-derived cells used in this study. P.S., T.E., L.M., and F.G. provided advice, supervised specific aspects of the work, and revised the manuscript. L.G. and L.V. wrote the manuscript.

## Extended Data Figures

**Extended Data Figure 1.**
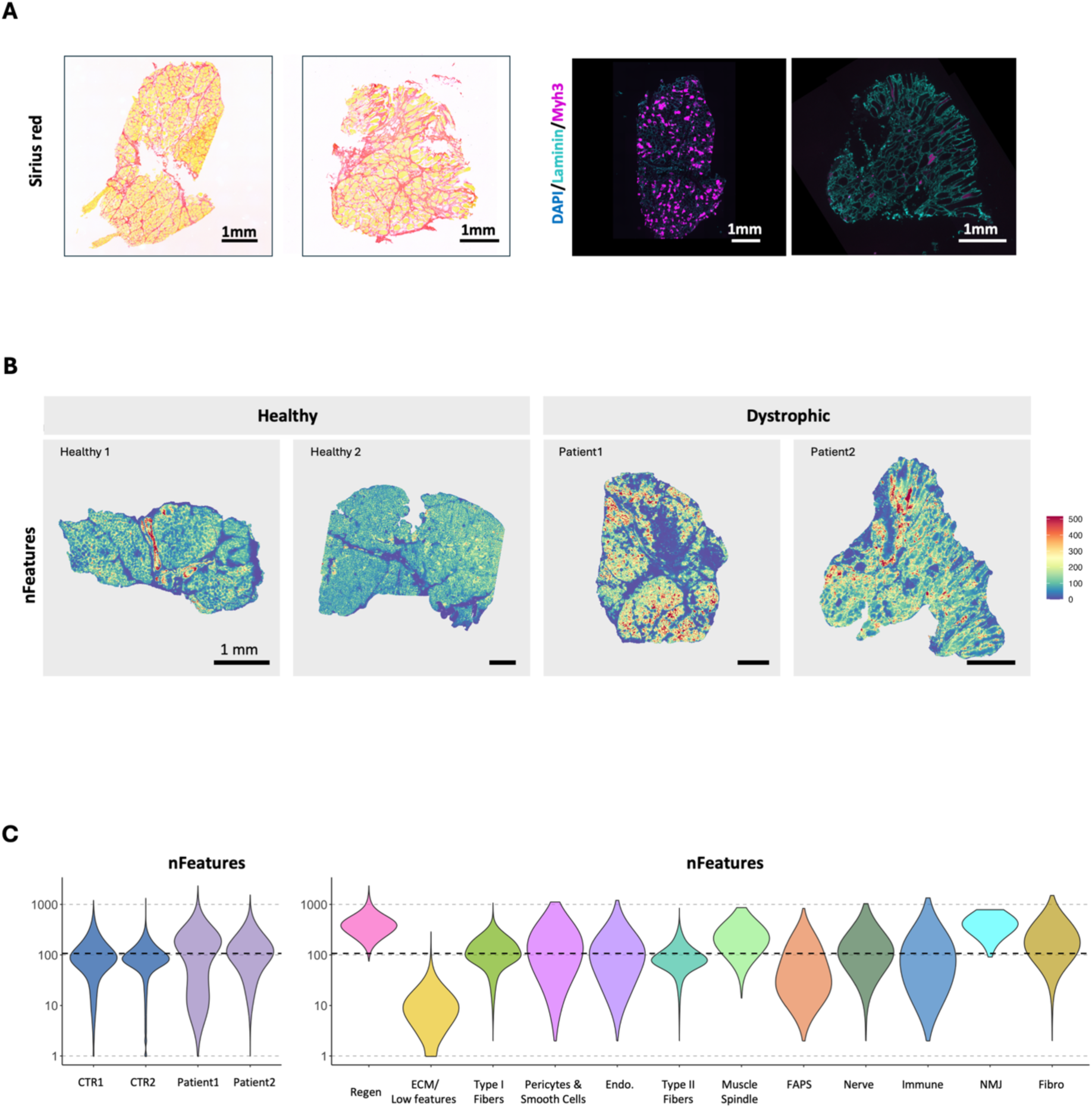
– Variability in transcript levels in different tissue regions and between samples. A) Sirius red (left) staining and immunostaining for Myh3 (Magenta), Laminin (Cyan) and Dapi (Blue) of DMD samples used for spatial transcriptomics (serial sections). Scale bar: 1mm. B) Spatial distribution of the number of features per spot across the different samples. Scale bar: 1 mm C) Violin plot displaying the number of features per sample (left) and per cluster (right).

**Extended Data Figure 2.**
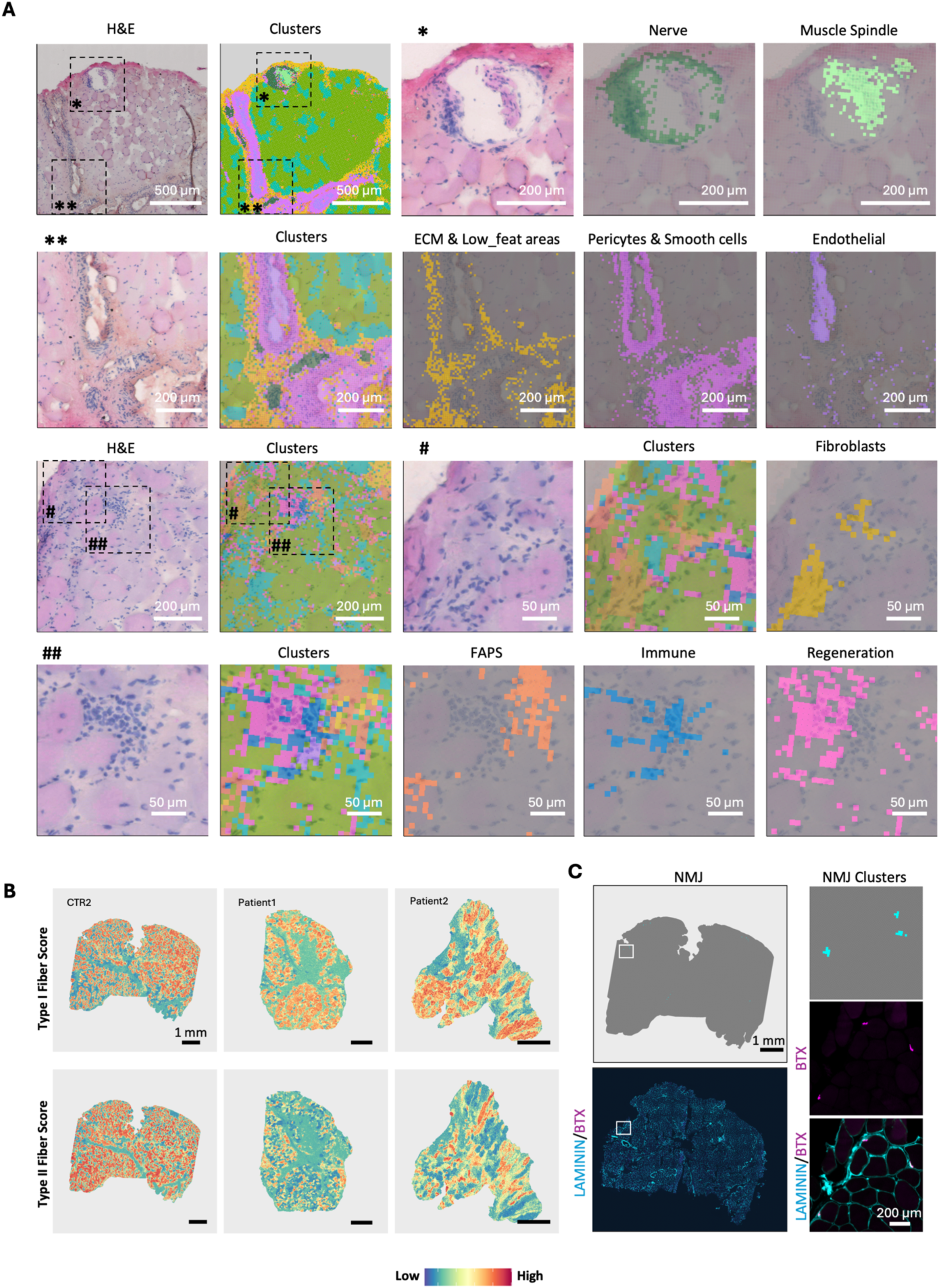
– Annotated cell populations correspond to anatomical structures observed in H&E-stained sections. A) Validation of selected cell population markers with known spatial localization. Scale bar specified in figure. The 8 µm bins are color-coded on the basis of their respective clusters. The symbols near the dashed box indicate the region shown in the inset. (B) Spatial distribution and relative expression levels of the fiber type score across different Visium HD samples. Scale bar: 1 mm. C) Overlap between the NMJ cluster-enriched region (top) and Bungarotoxin staining (bottom). A serial section was stained for Laminin (cyan) and Bungarotoxin (magenta) Scale bar: 1 mm. The white box indicates the region shown at higher magnification on the right Scale bar 200 µm.

**Extended Data Figure 3.**
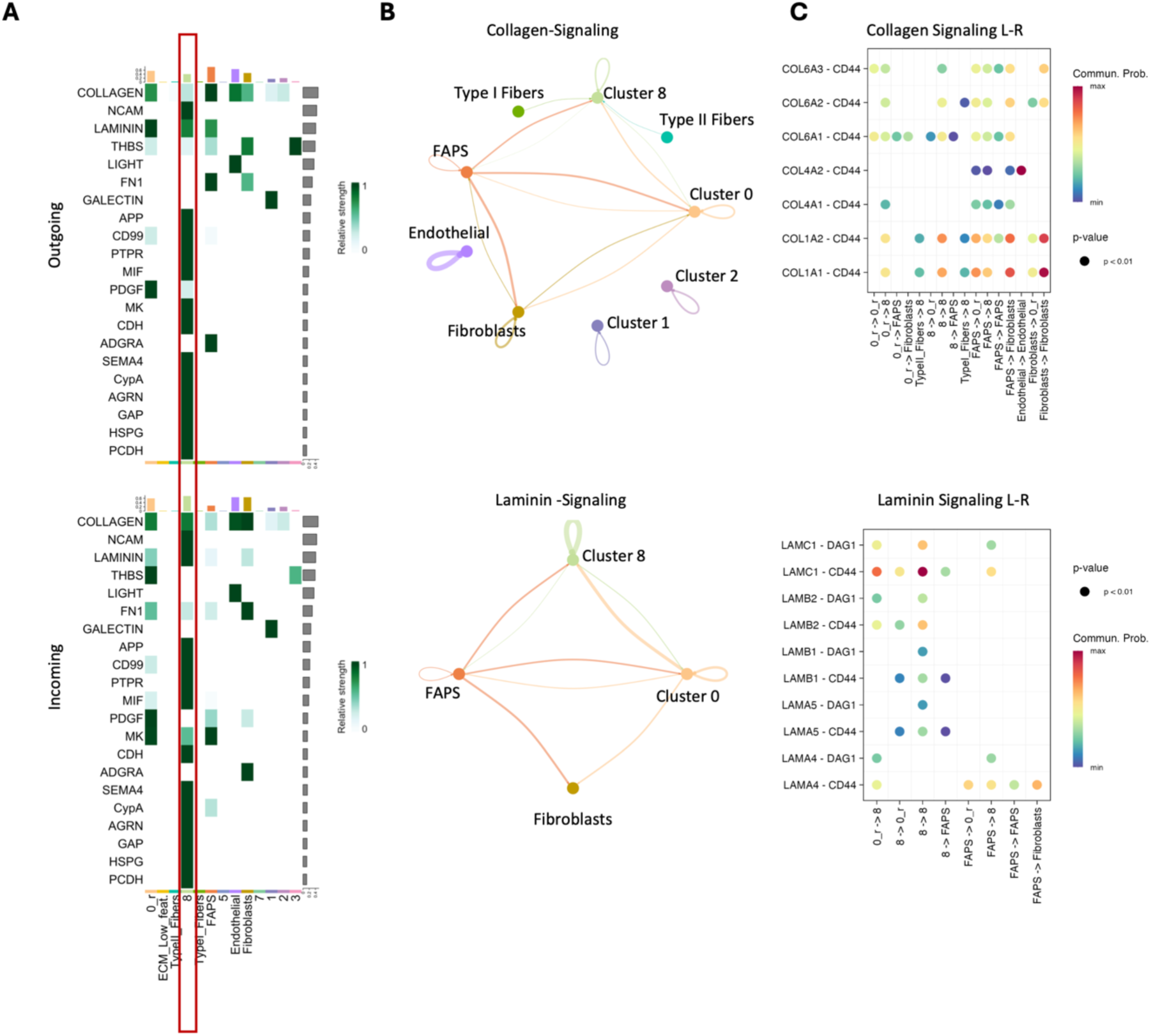
– Cell–cell communication analysis reveals ECM-driven signaling to regenerative clusters via CD44 in patient ST data. A) Heatmaps of the outgoing (top) and incoming (bottom) signals for the different clusters within the region. The color scale represents the relative interaction strength of each cell type for the represented pathways. B) Collagen (top) and Laminin (bottom) signaling networks. Nodes and edges are color-coded according to cell type. The edge width is proportional to the interaction probability. C) Dot plot representing the different ligand‒receptor pairs and the cell interaction pairs for Collagen (top) and Laminin signaling (bottom). The color scale represents the relative communication probability, and the dot size represents statistical significance.

**Extended Data Figure 4.**
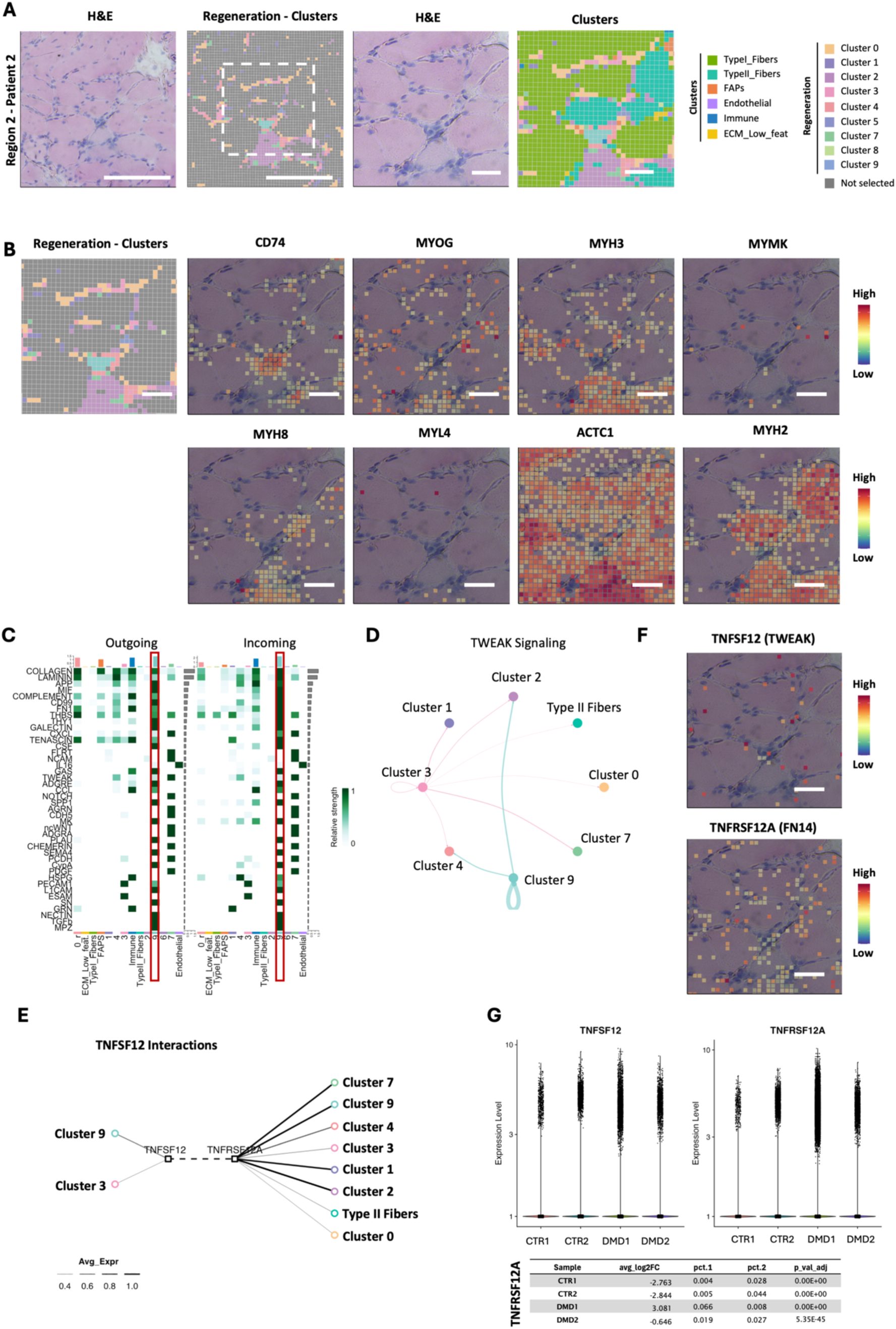
– Region 2 shows immune infiltration and TWEAK signaling in patient ST data. A) Selected region surrounding Clusters 2, 3 and 9: H&E staining and spatial distribution of regenerative subclusters (left) Scale bar: 200 µm. Zoomed-in regions showing H&E staining and combined spatial distribution of regeneration subclusters and cell type clusters (right) Scale bar: 50 µm. B) Spatial localization of regeneration subclusters and Expression of *CD74, MYOG, MYH3, MYMK, MYH8, MYL4, ACTC1,* and *MYH2* in region highlighted in (A) Scale bar: 50 µm. C) Heatmaps of the outgoing (left) and incoming (right) signals for the different clusters within the region. Color scale represents the relative interaction strength of each cell type for the represented pathways. D) TWEAK signaling network. Nodes and edges are color-coded according to the cell type. Edge width is proportional to the interaction probability. E) TNFSF12 mediated interactions and F) Expression of *TNFSF12* and *TNFRSF12A* within the regenerative area highlighted in (A). Scale bar: 50 µm G) Violin plot showing the relative expression levels for *TNFSF12* and *TNFRSF12A* across the samples (top). Table displaying average log₂ fold change, percentage of expressing bins (pct.1) within the sample vs. the rest of the dataset (pct.2), and statistical significance for *TNFRSF12A* (bottom). Statistical significance calculated using Wilcoxon rank-sum test as implemented in Seurat.

**Extended Data Figure 5.**
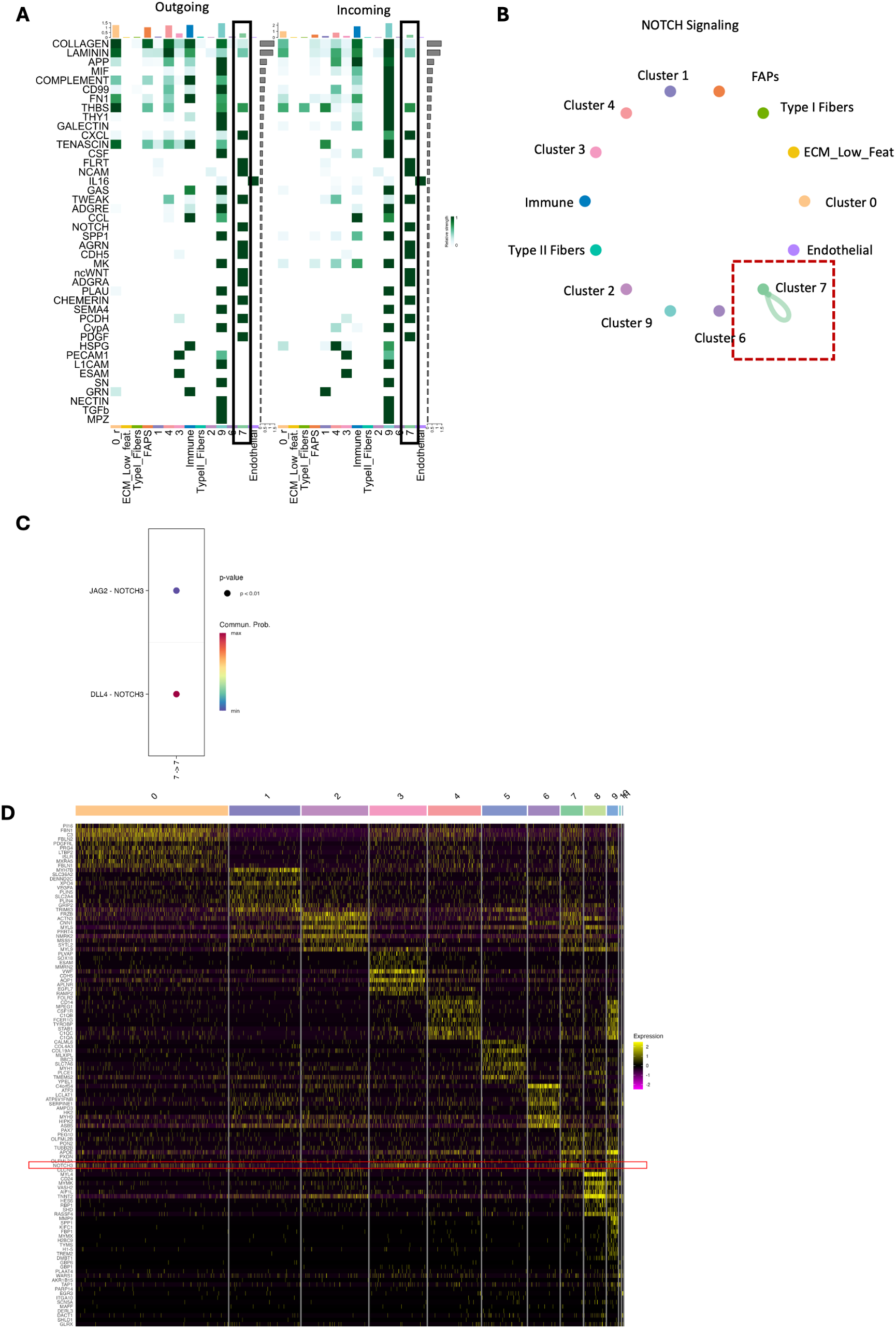
– Region 3 shows upregulated Notch3 signaling in patient ST data. A) Heatmaps of the outgoing (left) and incoming (right) signals for the different clusters within the highlighted area in Fig. 3D. The color scale represents the relative interaction strength of each cell type for the represented pathways. B) Notch signaling network within Region3. Nodes and edges are color-coded according to cell type. The edge width is proportional to the interaction probability. C) Dot plot representing the different ligand‒receptor pairs and the cell interaction pairs for Notch signaling. D) Heatmap showing expression values for the top 10 highest expressed genes for each regeneration subcluster after differential gene expression analysis. Genes were further filtered to include only those expressed in more than 10% of cells within the target population and in less than 20% of cells in the remaining populations.”

**Extended Data Figure 6.**
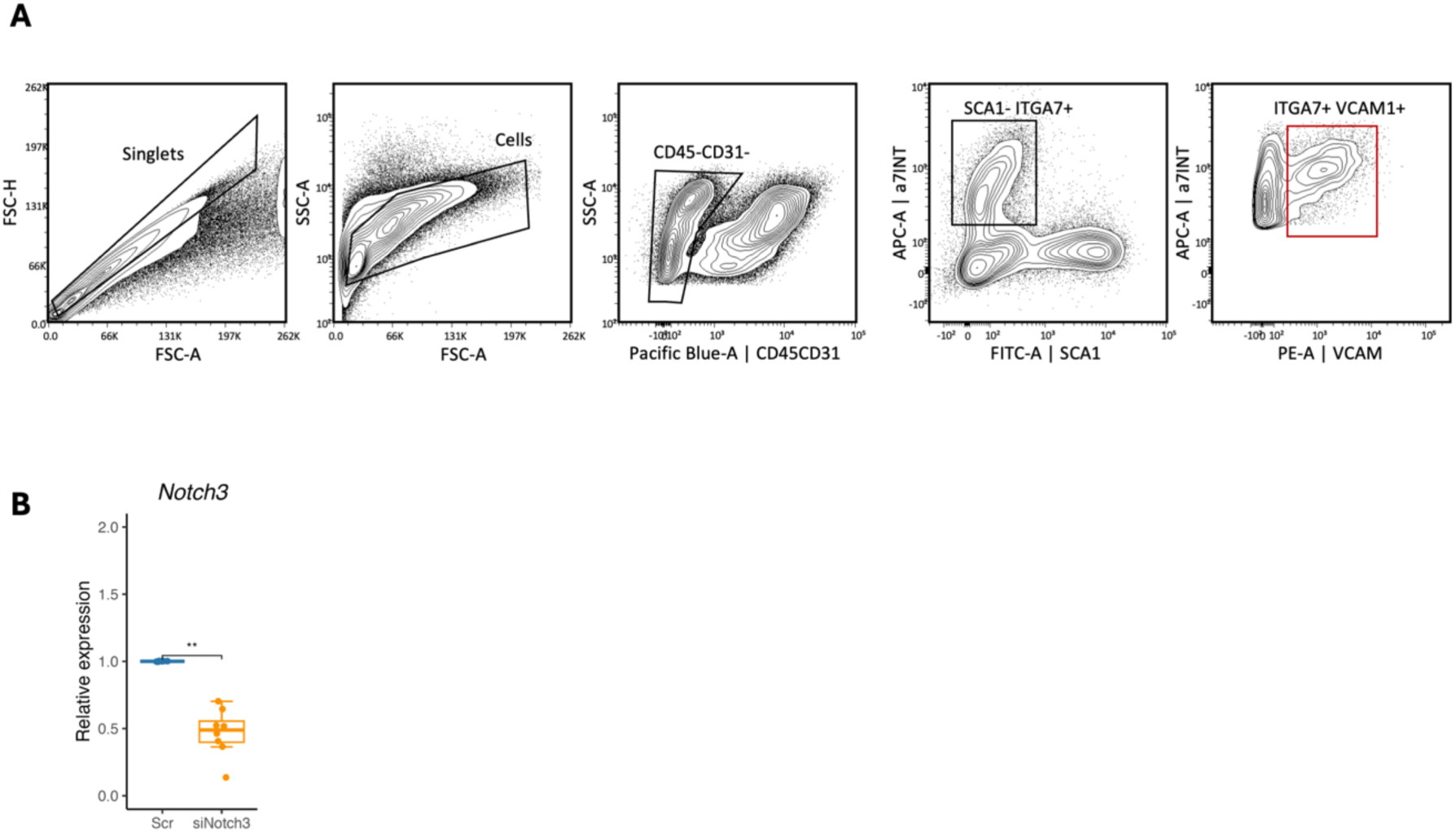
– MuSCs gating strategy and Notch3 levels after silencing in primary myoblasts. A) Flow plot illustrating the sorting strategy for MuSCs B) Relative expression of *Notch3*, in primary myoblasts from 6-month-old mdx mice at 48 h after siScr or siNotch3 transfection (n=8, paired Wilcoxon test). Boxplots show the 75th, 50th and 25th percentiles; whiskers extend to values within 1.5 × the interquartile range, * p < 0.05.

**Extended Data Figure 7.**
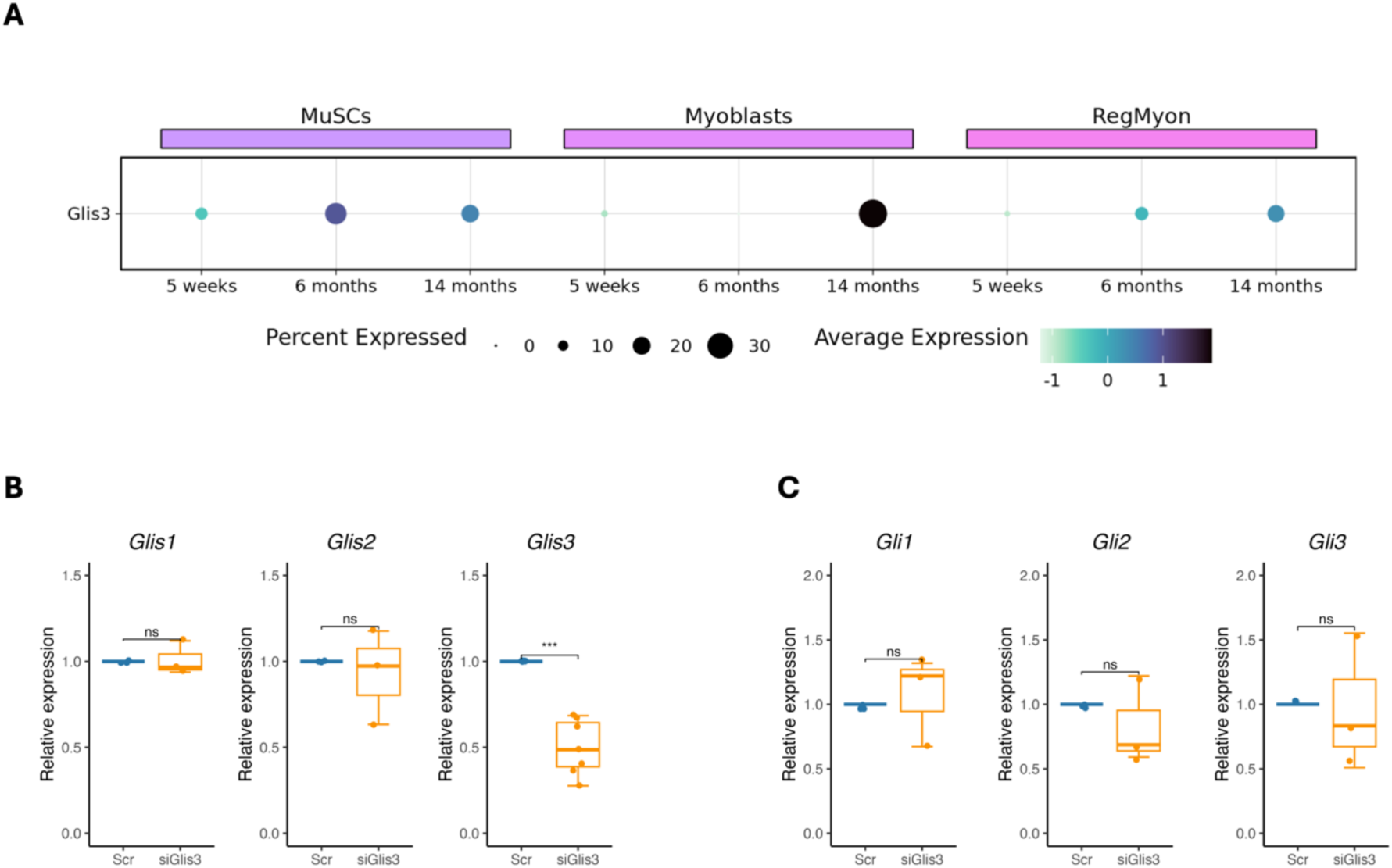
Glis3 is upregulated in multiple populations along myogenic trajectory at late stages of the disease. A) Dot plot showing the relative expression of *Glis3* across the different timepoints for the clusters within the myogenic trajectory. B) Relative expression of *Glis1* (n=3 – two-tailed paired t test), *Glis2* (n=3 – two-tailed paired t test) and *Glis3* (n=7– paired Wilcoxon test) in MuSCs from 6-month-old mdx mice at 48 h after siScr and siGlis3 transfection. C) Relative expression of *Gli1* (n=3), *Gli2* (n=3) and *Gli3* (n=3) in MuSCs from 6-month-old mdx mice at 48 h after siScr and siGlis3 transfection (two-tailed paired t test). Boxplots show the 75th, 50th and 25th percentiles; whiskers extend to values within 1.5 × the interquartile range, * p < 0.05.

**Extended Data Figure 8.**
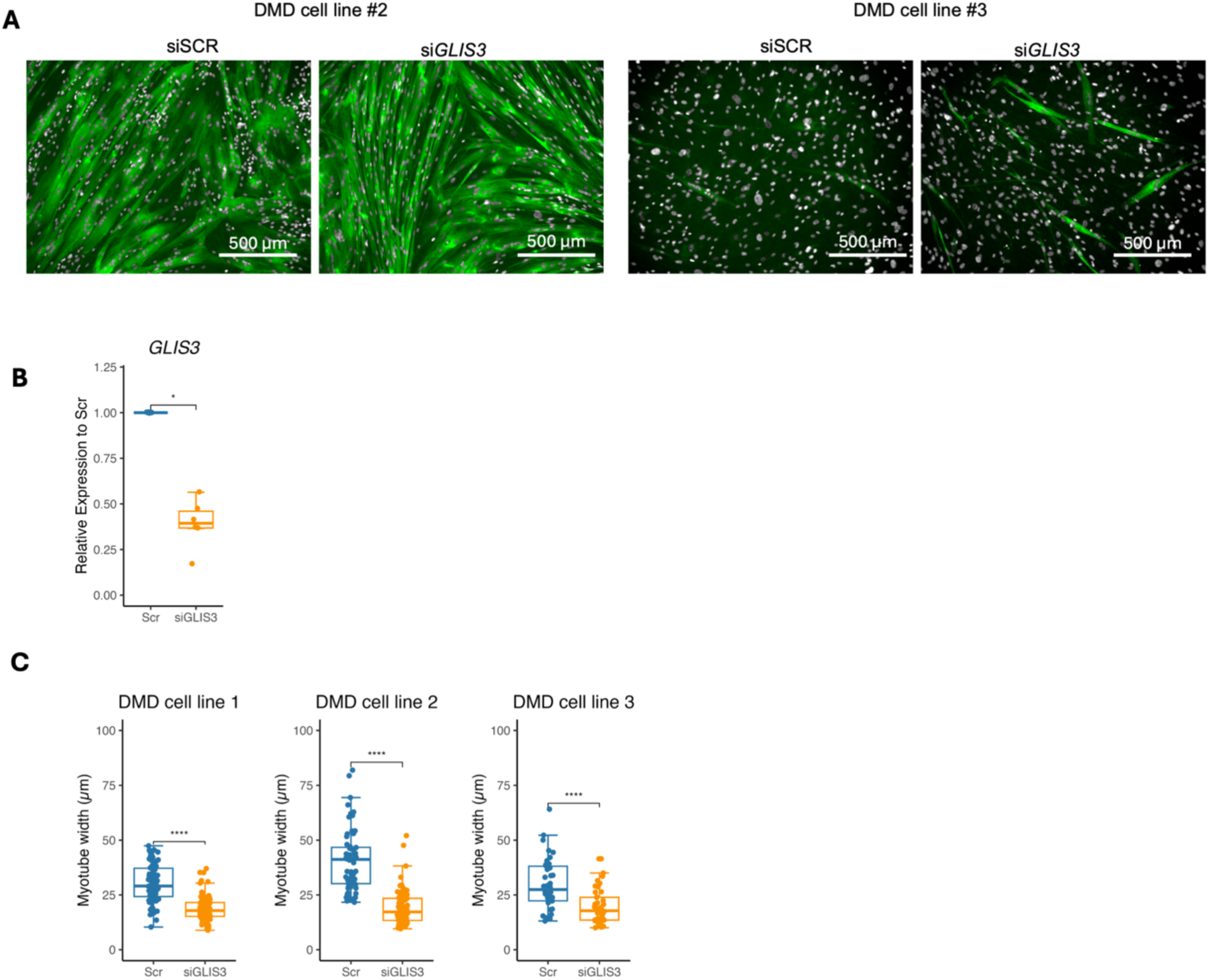
– Glis3 improves differentiation capacity but decreases myotube width in primary human cell lines. A) Representative images of human primary myoblast lines with different myogenic potentials transfected with siGLIS3 or nontargeting control siRNA (siSCR). Scale bar: 500 µm. B) Relative *GLIS3* levels in siGLIS3 and siSCR cells 48h post transfection (n= 6; 3 lines each with 2 independent technical replicates, Wilcoxon paired test. D) Myotube width quantification for three human cell lines with different myogenic potentials (at least 35 myotubes have been counted for each experimental point, two tailed Mann–Whitney U) Representative images of patient-derived lines are shown in A and in Fig. 6F, line #1. Boxplots show the 75th, 50th and 25th percentiles; whiskers extend to values within 1.5 × the interquartile range, * p < 0.05, ** p < 0.01, ***p < 0.001.

## Declaration of generative AI and AI-assisted technologies in the manuscript preparation process

During the preparation of this manuscript, the authors used ChatGPT (OpenAI) for spelling checks, grammar correction, and to improve the overall readability of the text. After using this tool/service, the author(s) reviewed and edited the content as needed and take(s) full responsibility for the content of the published article.

## References

1. Flanigan, K. M. Duchenne and Becker Muscular Dystrophies. Neurologic Clinics 32, 671–688 (2014).

2. Birnkrant, D. J. et al. Diagnosis and management of Duchenne muscular dystrophy, part 1: diagnosis, and neuromuscular, rehabilitation, endocrine, and gastrointestinal and nutritional management. Lancet Neurol 17, 251–267 (2018).

3. Dadgar, S. et al. Asynchronous remodeling is a driver of failed regeneration in Duchenne muscular dystrophy. J Cell Biol 207, 139–158 (2014).

4. Tidball, J. G., Welc, S. S. & Wehling-Henricks, M. Immunobiology of Inherited Muscular Dystrophies. Compr Physiol 8, 1313–1356 (2018).

5. Duan, D., Goemans, N., Takeda, S., Mercuri, E. & Aartsma-Rus, A. Duchenne muscular dystrophy. Nat Rev Dis Primers 7, 13 (2021).

6. Lemos, D. R. et al. Nilotinib reduces muscle fibrosis in chronic muscle injury by promoting TNF-mediated apoptosis of fibro/adipogenic progenitors. Nat. Med. 21, 786– 794 (2015).

7. Dumont, N. A. et al. Dystrophin expression in muscle stem cells regulates their polarity and asymmetric division. Nature Medicine 21, 1455–1463 (2015).

8. Chemello, F. et al. Degenerative and regenerative pathways underlying Duchenne muscular dystrophy revealed by single-nucleus RNA sequencing. PNAS 117, 29691–29701 (2020).

9. Tierney, M. T. et al. STAT3 signaling controls satellite cell expansion and skeletal muscle repair. Nat Med 20, 1182–1186 (2014).

10. Sarde, L. et al. Impaired stem cell migration and divisions in Duchenne Muscular Dystrophy revealed by live imaging. 2025.03.13.643016 Preprint at 10.1101/2025.03.13.643016 (2025).

11. Jiang, C. et al. Notch signaling deficiency underlies age-dependent depletion of satellite cells in muscular dystrophy. Dis Model Mech 7, 997–1004 (2014).

12. Church, J. E. et al. Alterations in Notch signalling in skeletal muscles from mdx and dko dystrophic mice and patients with Duchenne muscular dystrophy. Experimental Physiology 99, 675–687 (2014).

13. Saleh, K. K. et al. Single cell sequencing maps skeletal muscle cellular diversity as disease severity increases in dystrophic mouse models. iScience 25, 105415 (2022).

14. Scripture-Adams, D. D. et al. Single nuclei transcriptomics of muscle reveals intra-muscular cell dynamics linked to dystrophin loss and rescue. Commun Biol 5, 989 (2022).

15. Suárez-Calvet, X. et al. Decoding the transcriptome of Duchenne muscular dystrophy to the single nuclei level reveals clinical-genetic correlations. Cell Death Dis 14, 596 (2023).

16. Ren, S. et al. Profound cellular defects attribute to muscular pathogenesis in the rhesus monkey model of Duchenne muscular dystrophy. Cell 187, 6669–6686.e16 (2024).

17. Tierney, M. T. & Sacco, A. Inducing and Evaluating Skeletal Muscle Injury by Notexin and Barium Chloride. Methods Mol Biol 1460, 53–60 (2016).

18. Oliveira, M. F. et al. Characterization of immune cell populations in the tumor microenvironment of colorectal cancer using high definition spatial profiling. 2024.06.04.597233 Preprint at 10.1101/2024.06.04.597233 (2024).

19. Hao, Y. et al. Dictionary learning for integrative, multimodal and scalable single-cell analysis. Nat Biotechnol 42, 293–304 (2024).

20. McKellar, D. W. et al. Large-scale integration of single-cell transcriptomic data captures transitional progenitor states in mouse skeletal muscle regeneration. Commun Biol 4, 1–12 (2021).

21. D’Ercole, C. et al. Spatially resolved transcriptomics reveals innervation-responsive functional clusters in skeletal muscle. Cell Reports 41, 111861 (2022).

22. Heezen, L. G. M. et al. Spatial transcriptomics reveal markers of histopathological changes in Duchenne muscular dystrophy mouse models. Nat Commun 14, 4909 (2023).

23. Stec, M. J. et al. A cellular and molecular spatial atlas of dystrophic muscle. Proc Natl Acad Sci U S A 120, e2221249120.

24. Ruggieri, V. et al. Polyamine metabolism dysregulation contributes to muscle fiber vulnerability in ALS. Cell Reports 44, (2025).

25. Bornstein, B. et al. Molecular characterization of the intact mouse muscle spindle using a multi-omics approach. eLife 12, e81843.

26. Hicks, M. R. et al. Regenerating human skeletal muscle forms an emerging niche in vivo to support PAX7 cells. Nat Cell Biol 25, 1758–1773 (2023).

27. Lai, Y. et al. Multimodal cell atlas of the ageing human skeletal muscle. Nature 629, 154–164 (2024).

28. Murgia, M. et al. Protein profile of fiber types in human skeletal muscle: a single-fiber proteomics study. Skeletal Muscle 11, 24 (2021).

29. Chaillou, T. et al. Identification of a conserved set of upregulated genes in mouse skeletal muscle hypertrophy and regrowth. J Appl Physiol (1985) 118, 86–97 (2015).

30. Urciuolo, A. et al. Collagen VI regulates satellite cell self-renewal and muscle regeneration. Nat Commun 4, 1964 (2013).

31. Lehto, M., Duance, V. C. & Restall, D. Collagen and fibronectin in a healing skeletal muscle injury. An immunohistological study of the effects of physical activity on the repair of injured gastrocnemius muscle in the rat. J Bone Joint Surg Br 67, 820–828 (1985).

32. Fitzgerald, G. et al. MME+ fibro-adipogenic progenitors are the dominant adipogenic population during fatty infiltration in human skeletal muscle. Commun Biol 6, 1–21 (2023).

33. Jin, S., Plikus, M. V. & Nie, Q. CellChat for systematic analysis of cell–cell communication from single-cell transcriptomics. Nat Protoc 20, 180–219 (2025).

34. Mylona, E., Jones, K. A., Mills, S. T. & Pavlath, G. K. CD44 regulates myoblast migration and differentiation. Journal of Cellular Physiology 209, 314–321 (2006).

35. Bentzinger, C. F. et al. Fibronectin regulates Wnt7a signaling and satellite cell expansion. Cell Stem Cell 12, 75–87 (2013).

36. Girgenrath, M. et al. TWEAK, via its receptor Fn14, is a novel regulator of mesenchymal progenitor cells and skeletal muscle regeneration. EMBO J 25, 5826–5839 (2006).

37. Morosetti, R. et al. TWEAK in Inclusion-Body Myositis Muscle: Possible Pathogenic Role of a Cytokine Inhibiting Myogenesis. The American Journal of Pathology 180, 1603–1613 (2012).

38. Dogra, C., Hall, S. L., Wedhas, N., Linkhart, T. A. & Kumar, A. Fibroblast growth factor inducible-14 (Fn14) is required for the expression of myogenic regulatory factors and differentiation of myoblasts into myotubes: Evidence for TWEAK-independent functions of Fn14 during myogenesis. J Biol Chem 282, 15000–15010 (2007).

39. Conboy, I. M., Conboy, M. J., Smythe, G. M. & Rando, T. A. Notch-Mediated Restoration of Regenerative Potential to Aged Muscle. Science 302, 1575–1577 (2003).

40. Fukada, S. et al. Molecular signature of quiescent satellite cells in adult skeletal muscle. Stem Cells 25, 2448–2459 (2007).

41. Kuang, S., Kuroda, K., Le Grand, F. & Rudnicki, M. A. Asymmetric self-renewal and commitment of satellite stem cells in muscle. Cell 129, 999–1010 (2007).

42. Bjornson, C. R. R. et al. Notch signaling is necessary to maintain quiescence in adult muscle stem cells. Stem Cells 30, 232–42 (2012).

43. Kitamoto, T. & Hanaoka, K. Notch3 Null Mutation in Mice Causes Muscle Hyperplasia by Repetitive Muscle Regeneration. Stem Cells 28, 2205–2216 (2010).

44. Mourikis, P. et al. A Critical Requirement for Notch Signaling in Maintenance of the Quiescent Skeletal Muscle Stem Cell State. Stem Cells 30, 243–252 (2012).

45. Baghdadi, M. B. et al. Reciprocal signalling by Notch–Collagen V–CALCR retains muscle stem cells in their niche. Nature 557, 714–718 (2018).

46. Verma, M. et al. Muscle Satellite Cell Cross-Talk with a Vascular Niche Maintains Quiescence via VEGF and Notch Signaling. Cell Stem Cell 23, 530–543.e9 (2018).

47. Lahmann, I. et al. Oscillations of MyoD and Hes1 proteins regulate the maintenance of activated muscle stem cells. Genes Dev. 33, 524–535 (2019).

48. Yartseva, V. et al. Heterogeneity of Satellite Cells Implicates DELTA1/NOTCH2 Signaling in Self-Renewal. Cell Reports 30, 1491–1503.e6 (2020).

49. Zhang, Y. et al. Oscillations of Delta-like1 regulate the balance between differentiation and maintenance of muscle stem cells. Nat Commun 12, 1318 (2021).

50. Jiang, C. et al. Notch signaling deficiency underlies age-dependent depletion of satellite cells in muscular dystrophy. Dis Model Mech 7, 997–1004 (2014).

51. Giordani, L. et al. High-Dimensional Single-Cell Cartography Reveals Novel Skeletal Muscle-Resident Cell Populations. Molecular Cell 74, 609–621.e6 (2019).

52. De Micheli, A. J., Spector, J. A., Elemento, O. & Cosgrove, B. D. A reference single-cell transcriptomic atlas of human skeletal muscle tissue reveals bifurcated muscle stem cell populations. Skeletal Muscle 10, 19 (2020).

53. Dell’Orso, S. et al. Single cell analysis of adult mouse skeletal muscle stem cells in homeostatic and regenerative conditions. Development 146, (2019).

54. Dos Santos, M. et al. Single-nucleus RNA-seq and FISH identify coordinated transcriptional activity in mammalian myofibers. Nature Communications 11, 5102 (2020).

55. Petrany, M. J. et al. Single-nucleus RNA-seq identifies transcriptional heterogeneity in multinucleated skeletal myofibers. Nature Communications 11, 6374 (2020).

56. Kim, M. et al. Single-nucleus transcriptomics reveals functional compartmentalization in syncytial skeletal muscle cells. Nature Communications 11, 6375 (2020).

57. Irintchev, A., Zweyer, M. & Wernig, A. Impaired functional and structural recovery after muscle injury in dystrophic *mdx* mice. Neuromuscular Disorders 7, 117–125 (1997).

58. Pastoret, C. & Sebille, A. *mdx* mice show progressive weakness and muscle deterioration with age. Journal of the Neurological Sciences 129, 97–105 (1995).

59. Low, S., Barnes, J. L., Zammit, P. S. & Beauchamp, J. R. Delta-Like 4 Activates Notch 3 to Regulate Self-Renewal in Skeletal Muscle Stem Cells. Stem Cells 36, 458–466 (2018).

60. Kamimoto, K. et al. Dissecting cell identity via network inference and in silico gene perturbation. Nature 1–10 (2023) doi:10.1038/s41586-022-05688-9.

61. Schep, A. N., Wu, B., Buenrostro, J. D. & Greenleaf, W. J. chromVAR: inferring transcription-factor-associated accessibility from single-cell epigenomic data. Nat Methods 14, 975–978 (2017).

62. Kim, Y.-S., Nakanishi, G., Lewandoski, M. & Jetten, A. M. GLIS3, a novel member of the GLIS subfamily of Krüppel-like zinc finger proteins with repressor and activation functions. Nucleic Acids Res 31, 5513–5525 (2003).

63. Scoville, D. W., Kang, H. S. & Jetten, A. M. GLIS1-3: emerging roles in reprogramming, stem and progenitor cell differentiation and maintenance. Stem Cell Investig 4, 80 (2017).

64. Jetten, A. M. GLIS1–3 transcription factors: critical roles in the regulation of multiple physiological processes and diseases. Cell Mol Life Sci 75, 3473–3494 (2018).

65. Duan, H., Skeath, J. B. & Nguyen, H. T. Drosophila Lame duck, a novel member of the Gli superfamily, acts as a key regulator of myogenesis by controlling fusion-competent myoblast development. Development 128, 4489–4500 (2001).

66. Duan, H. & Nguyen, H. T. Distinct Posttranscriptional Mechanisms Regulate the Activity of the Zn Finger Transcription Factor Lame duck during Drosophila Myogenesis. Mol Cell Biol 26, 1414–1423 (2006).

67. Yadava, R. S. et al. TWEAK Regulates Muscle Functions in a Mouse Model of RNA Toxicity. PLoS One 11, e0150192 (2016).

68. Yadava, R. S. et al. TWEAK/Fn14, a pathway and novel therapeutic target in myotonic dystrophy. Hum Mol Genet 24, 2035–2048 (2015).

69. Pescatori, M. et al. Gene expression profiling in the early phases of DMD: a constant molecular signature characterizes DMD muscle from early postnatal life throughout disease progression. The FASEB Journal 21, 1210–1226 (2007).

70. Burkly, L. C., Michaelson, J. S. & Zheng, T. S. TWEAK/Fn14 pathway: an immunological switch for shaping tissue responses. Immunological Reviews 244, 99–114 (2011).

71. Kang, H. S. et al. Transcription Factor GLIS3: A New and Critical Regulator of Postnatal Stages of Mouse Spermatogenesis. Stem Cells 34, 2772–2783 (2016).

72. Aloisio, G. M. et al. PAX7 expression defines germline stem cells in the adult testis. J Clin Invest 124, 3929–3944 (2014).

73. Tosic, M. et al. Lsd1 regulates skeletal muscle regeneration and directs the fate of satellite cells. Nat Commun 9, 366 (2018).

74. Brun, C. E. et al. GLI3 regulates muscle stem cell entry into GAlert and self-renewal. Nat Commun 13, 3961 (2022).

75. Vissing, K. & Schjerling, P. Simplified data access on human skeletal muscle transcriptome responses to differentiated exercise. Sci Data 1, 140041 (2014).

76. Bulfield, G., Siller, W. G., Wight, P. A. & Moore, K. J. X chromosome-linked muscular dystrophy (mdx) in the mouse. Proceedings of the National Academy of Sciences 81, 1189–1192 (1984).

77. Liu, L., Cheung, T. H., Charville, G. W. & Rando, T. A. Isolation of skeletal muscle stem cells by fluorescence-activated cell sorting. Nat. Protocols 10, 1612–1624 (2015).

78. Wickham, H. Ggplot2: Elegant Graphics for Data Analysis. (Springer-Verlag New York, 2016).

79. Wickham H, François R, Henry L, Müller K, Vaughan D. dplyr: A Grammar of Data Manipulation. R package version 1.1.4 https://dplyr.tidyverse.org/.

80. Pedersen, T. L. Patchwork: The Composer of Plots. (2025).

81. Stuart, T., Srivastava, A., Madad, S., Lareau, C. & Satija, R. Single-cell chromatin state analysis with Signac. Nature Methods 10.1038/s41592-021-01282-5 (2021) doi:10.1038/s41592-021-01282-5.

82. Gu, Z., Eils, R. & Schlesner, M. Complex heatmaps reveal patterns and correlations in multidimensional genomic data. Bioinformatics 32, 2847–2849 (2016).

83. Csárdi, G. & Nepusz, T. The igraph software package for complex network research. InterJournal Complex Systems, 1695 (2006).

84. Rainer, J., Gatto, L. & Weichenberger, C. X. ensembldb: an R package to create and use Ensembl-based annotation resources. Bioinformatics 10.1093/bioinformatics/btz031 (2019) doi:10.1093/bioinformatics/btz031.

85. Lawrence, M. et al. Software for Computing and Annotating Genomic Ranges. PLOS Computational Biology 9, e1003118 (2013).

